# A universal taxonomic and functional human gut microbiome model for disease classification and phenotype discovery

**DOI:** 10.64898/2026.04.30.721924

**Authors:** Zuzanna Karwowska, Marcin Możejko, Wojciech Nowak, Anastasia Romanchenko, Ewa Szczurek, Tomasz Kosciolek

## Abstract

The human gut microbiome is a powerful indicator of host health, yet its compositional nature, high sparsity, and inter-individual variability complicate downstream analysis. Here, we introduce two complementary approaches to characterize gut microbiome structure at population scale. First, we define eight functional signatures of the human gut microbiome using Non-negative Matrix Factorization, revealing coordinated metabolic patterns that partially decouple from taxonomic composition. Second, we present GUT-FORMer, a transformer-based autoencoder that jointly models taxonomic and functional metagenomic profiles from close to 21,000 publicly available samples. The learned latent representations capture biologically meaningful structure, reflect geographic and disease-associated variation, and enable accurate classification of 25 diseases in both binary and multiclass settings, as well as regression of host age and BMI. GUT-FORMer outperforms existing microbiome indices and deep learning methods across all tasks, establishing a generalizable framework for microbiome-based precision medicine.

## INTRODUCTION

The connection between the human gut microbiome and host health has been extensively studied over the past decades (Asnicar et al. 2026; Corral López et al. 2026; Ferretti et al. 2026). The gut microbiome contributes to digestion (Fu et al. 2022), vitamin production(Tarracchini et al. 2024), immune regulation(Zheng et al. 2020), host metabolism (Jyoti and Dey 2025), and neurological processes (Doenyas et al. 2025), and is increasingly explored as a therapeutic target through approaches such as dietary interventions (Asnicar et al. 2026), probiotics (Sanders et al. 2019), fecal microbiota transplantation (Ianiro et al. 2022), and live biotherapeutics (Pribyl et al. 2025).

Despite its importance, the gut microbiome remains challenging to study due to both biological and computational factors (Kumar et al. 2024). Microbiome sequencing data are compositional, meaning abundances are relative and constrained by sequencing depth, and highly sparse, as individuals share relatively few microbial species (Gloor et al. 2017). In addition, microbiome composition varies substantially across individuals and populations due to factors such as diet, lifestyle, geography, medication use, and temporal dynamics (Vujkovic-Cvijin et al. 2020; Aasmets et al. 2025). These properties complicate tasks such as identifying disease-associated microbial signatures, predicting therapeutic responses, or designing microbiome-based interventions.

Several approaches have attempted to identify structure in gut microbiome data, including enterotypes (Arumugam et al. 2011), enterosignatures (Frioux et al. 2023), and more recent machine learning methods that learn latent representations of microbial communities (Zahavi et al. 2025; Zhang et al. 2026; Pope et al. 2025). However, many existing approaches rely primarily on taxonomic composition, which describes which organisms are present but provides limited insight into the functional capabilities of the microbiome (Zielińska et al. 2025).

Metagenomic functional profiling offers a complementary perspective by capturing the metabolic potential of microbial communities (Peng et al. 2025; Manghi et al. 2025). Functional data are often less sparse than taxonomic profiles, and individuals that differ taxonomically may nevertheless share similar functional potential (Zielińska et al. 2025; Maranga et al. 2023). However, functional information alone is also incomplete, as many microbial genes remain unannotated and functions are ultimately carried by specific organisms (Maranga et al. 2023).

To address these challenges, we introduce two complementary approaches. First, we define functional signatures of the human gut microbiome, representing characteristic combinations of metabolic pathways that capture population-level variation in microbiome function. Second, we present GUT-FORMer (Gut microbiome Universal Taxonomy and Function Transformer), a deep learning framework that jointly models taxonomic composition and functional potential using a transformer-based autoencoder architecture with a loss function adapted to compositional microbiome data (Figure 1).

**Figure 1.**
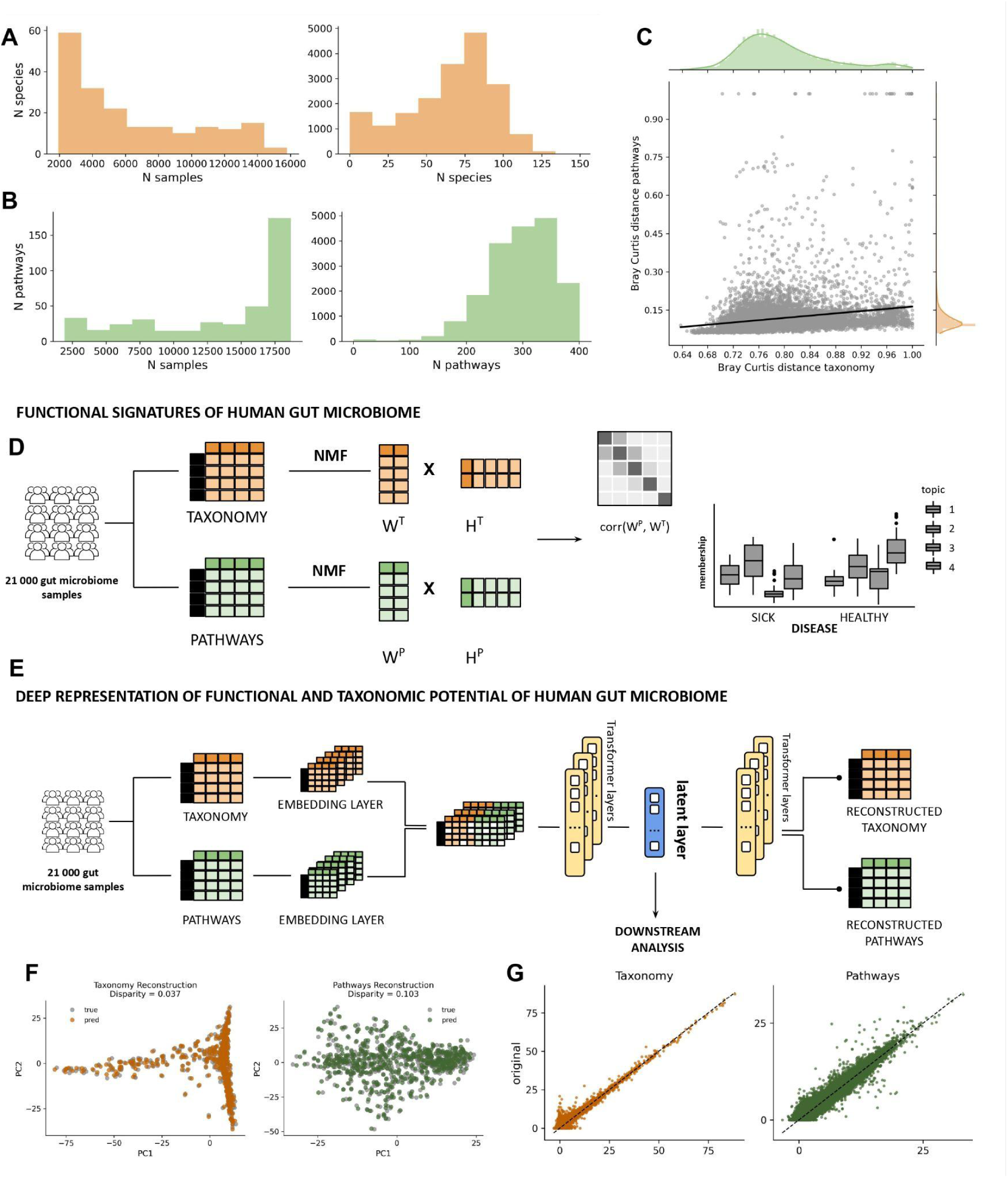
Data and models overview. A. Distribution of unique species across samples; B. Distribution of unique pathways across samples; C. Comparison of the mean Bray-Curtis distance between subjects calculated on taxonomy and pathways; D. Signatures of the human gut microbiome obtained by fitting Non-Negative matrix factorization (NMF). NMF is fitted on taxonomy which creates matrices W - abundance of specific topics across subjects and H - feature contribution to specific topics. Using the same approach functional signatures of the human gut microbiome are calculated. Obtained representation using NMF can be then used to understand relationship between pathways and taxonomy in the human gut microbiome and in order to better understand specific patterns in health and disease. E. GUT-FORMer auto encoder model with transformer layers trained on taxonomy and pathways to learn comprehensive and biology-driven representation of the human gut microbiome. F. Sample-wise comparison of input and reconstructed taxonomy and functional data using Procrustes. Disparity represents .. G. Feature-wise comparison between input and reconstructed taxonomy and pathways matrices.

We show that GUT-FORMer learns biologically meaningful representations of microbial communities, enabling improved exploration of microbiome structure and accurate classification of host phenotypes, including disease status, as well as regression tasks: age, and BMI prediction. Moreover, the learned representations outperform several existing microbiome health indices and previously proposed deep learning models.

## RESULTS

### MOTIVATION AND MODEL OVERVIEW

Gut microbiome taxonomic feature tables are highly sparse ranging from 75-90% of zeros(Busato et al. 2023). Many bacterial species are present in only a small subset of individuals, with limited overlap between subjects (Fig 1A). High sparsity and inter-individual variability make taxonomic analysis challenging and complicates biomarker discovery, representation learning, differential abundance analysis, and estimation of appropriate sample sizes (Nearing et al. 2022) .

Functional metagenomic data (e.g., metabolic pathways) is much less sparse (Fig. 1B) as most functional pathways are present in the majority of individuals (Zielińska et al. 2025).

Functional metagenomic data provide information complementary to taxonomic composition - individuals can differ substantially at the taxonomic level while remaining similar at the functional level (Fig 1C). This might suggest that taxonomically distinct microbial communities may perform similar biological functions and that functional annotation helps reveal biologically meaningful similarities and differences that may not be apparent from taxonomy alone.

### FUNCTIONAL SIGNATURES OF HUMAN GUT MICROBIOME

Motivated by the fact that functional metagenomic data is less sparse and the concept of taxonomic enterosignatures introduced by Frioux et al. (Frioux et al. 2023), we introduce functional signatures of the human gut microbiome (Fig. 1D). Using the Non-negative matrix factorisation (NMF)-based approach we analyze if functional signatures are related to taxonomic features and whether they can be used to understand specific patterns in gut microbiomes linked to health and disease. Further, we show that functional signatures are strongly associated with taxonomic signatures, indicating that functional organization is linked to underlying microbial composition. Finally, we demonstrate that functional signatures show improved performance in distinguishing healthy and diseased individuals compared to taxonomic signatures alone. We believe that functional signatures may provide more robust and biologically meaningful features for microbiome-based classification and biomarker discovery.

### GUT-FORMer MODEL

Taxonomic and functional microbiome data are both necessary and complementary. Functional data provides insight into the biological activity and metabolic potential of the microbiome, revealing what functions are being performed, while taxonomic data identify the microbial organisms responsible for these functions, enabling attribution of functional activity to specific species. Integrating both data types is essential for a complete and interpretable understanding of gut microbiome structure and function.

Motivated by the strong signal captured by NMF-based functional and taxonomy-level signatures, we developed a deep learning model, GUT-FORMer, which leverages both taxonomic and functional information derived from metagenomic samples.

GUT-FORMer is a transformer-based autoencoder architecture trained jointly on taxonomic and functional profiles derived from the same individuals. The model was trained on publicly available human gut metagenomic samples from the CuratedMetagenomicData resource (Pasolli et al. 2017). The training dataset encompasses samples from healthy controls and individuals with various diseases, from different countries across the world, across age groups ranging from newborns to seniors, and is evenly distributed across sexes (supplementary Figure 1A-D). The objective of GUT-FORMer is to learn an unified representation of the human gut microbiome that integrates both taxonomic composition and its functional potential, enabling the identification of latent patterns and structure across populations (Figure 1E).

We demonstrate that GUT-FORMer accurately reconstructs microbiome profiles, both: subject level (overall microbiome structure), and at the feature level (individual taxa and functional pathways) (Figure 1F,G). The learned latent representations are biologically interpretable and capture meaningful microbiome structure consistent with NMF-derived signatures at both taxonomic and functional levels. We further test if the latent representation provides informative features for downstream tasks, including classification and regression and if GUT-FORMer outperforms existing methods in downstream predictive tasks including classification and regression.

### FUNCTIONAL SIGNATURES OF THE HUMAN GUT MICROBIOME

Using an approach based on nine-fold bicross-validation was performed as described in Owen and Perry (Owen and Perry 2009), Kanagal and Sindhwani (Kanagal and Sindhwani 2010), and Eng et al. (Eng et al. 2019) we propose 8 functional signatures of the human gut microbiome.

Each signature can be defined by a different set of functions (Fig. 2A) but when compared to taxonomic signatures, functional topics are much more diverse with multiple features influencing each signature. A full list of functions that contribute to each signature can be found in supplementary Figure 2.

**Figure 2.**
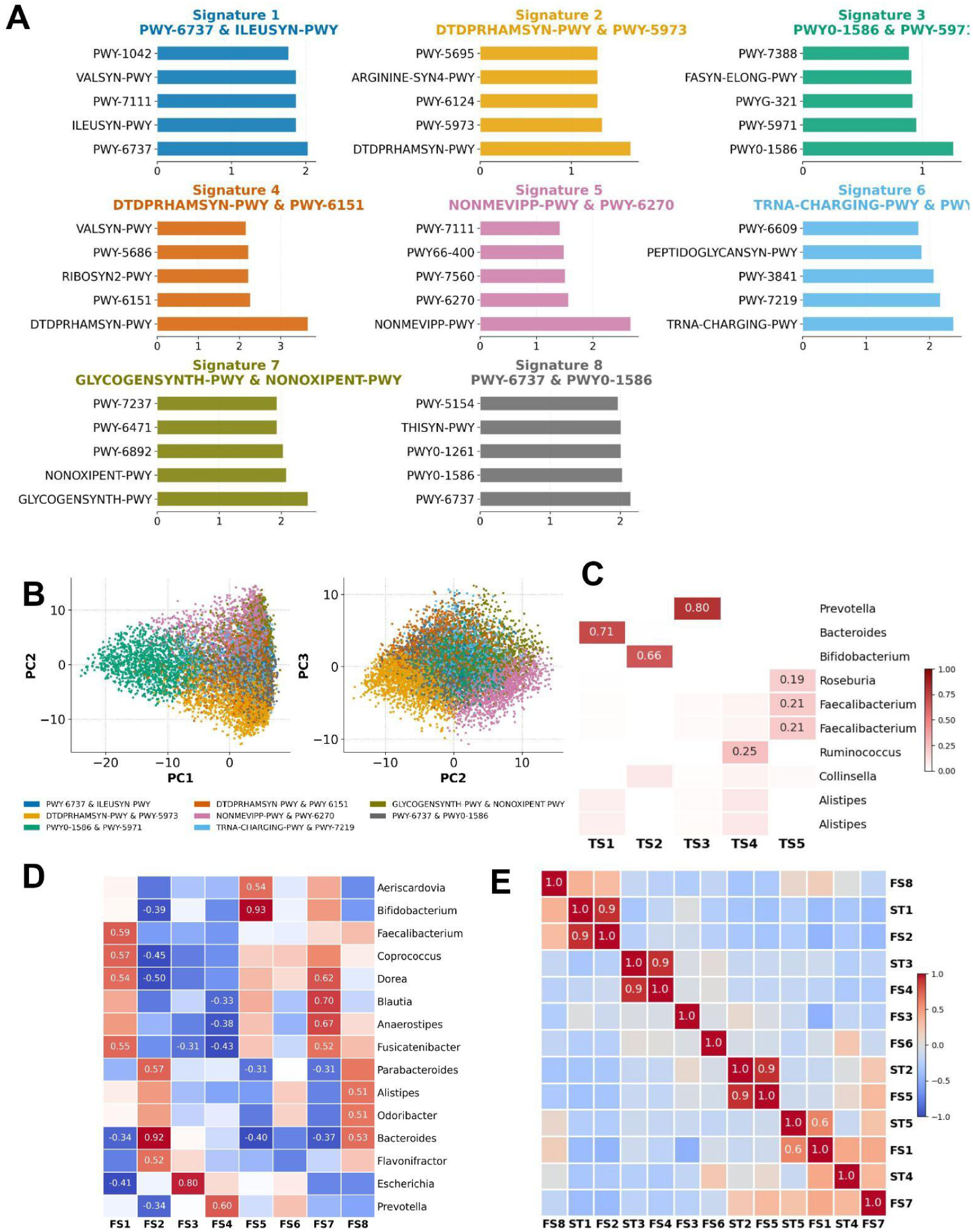
Functional signatures of the human gut microbiome. A. Features contribution to each topic. Top 5 features are shown. B. PCoA on pathways using Atchison distance, subjects are colored by the dominant functional signature. C. Taxonomy-based signatures obtained by NMF (Frioux et al.). D. Pearson correlation between genus and functional signatures. E. Pearson correlation between functional and taxonomic signatures of the human gut microbiome. Abbreviations: FS - functional signature; TS - taxonomic signature

PCoA (Principal Coordinate Analysis) of functional abundance shows that functional signatures differentiate subjects. In particular, signatures 2 and 5 contribute most strongly to the separation as subjects dominated by these signatures are the most distinct in the ordination space, suggesting distinct functional patterns among these individuals (Fig. 2B). Compared to the taxonomic enterosignatures introduced by Frioux et al., functional signatures appear less driven by individual features: each signature is present across many subjects at varying abundances. This suggests that, unlike taxonomic signatures—which are often dominated by a single genus—functional profiles are driven by an ensemble of microbial functions rather than by individual taxa. Signature 1 is present in most of the samples which might suggest a core metabolic signature (supplementary Figure 2B).

Analysis of the distribution of functional signatures across diseases further reveals disease-specific patterns. While signatures 1 and 2 are the most abundant across subjects regardless of disease, other signatures vary more substantially between conditions. This may indicate that signatures 1 and 2 represent core functional modules of the gut microbiome that are broadly conserved across individuals, whereas the remaining signatures capture functional shifts associated with particular disease states (Supplementary Figure 2C).

Pearson correlation analysis revealed clear associations between functional signatures and taxonomic composition at both the genus and enterosignature levels (Fig. 2C–E).

At the genus level (Fig. 2D), functional signatures exhibited varying degrees of specificity. Signatures 1 and 8 were broadly associated with multiple genera, whereas others showed more targeted relationships, including Signature 2 with *Bacteroides*, Signature 3 with *Escherichia*, Signature 4 with *Prevotella*, Signature 5 with *Bifidobacterium*, Signature 6 with *Prevotella* and Signature 7 with *Blautia*, *Anaerostipes*, and *Dorea*. In contrast, Signature 4 did not display strong associations with any single genus, suggesting it may represent functional modules distributed across multiple taxa.

Consistent patterns were observed at the enterosignature level (Fig. 2D-E), based on reference signatures defined in healthy individuals (Frioux et al.). Most functional signatures align well with taxonomic enterosignatures, reinforcing the link between microbial composition and functional potential. However, Signatures 3, 6, and 8 showed weak or no associations, supporting the idea that some functional modules are not restricted to specific taxonomic groups but are shared across diverse microbial lineages.

### GUT-FORMER LATENT SPACE CAPTURES BIOLOGICALLY RELEVANT STRUCTURE IN THE HUMAN GUT MICROBIOME

#### Latent representation

We used PHATE (Potential of Heat-diffusion for Affinity-based Trajectory Embedding) to visualize the latent space, revealing clear and structured organization of microbiome samples (Moon et al. 2019)). PHATE is a visualization method that captures both local and global nonlinear structure in data by an information-geometric distance between datapoints . We ran PHATE with default parameters.

Previously defined taxonomic signatures are clearly reflected in the latent representation (Fig. 3A). Functional signatures defined in Fig. 2 are also clearly captured in the latent space (Fig. 3B). The representation demonstrates that taxonomic and functional signatures are complementary: some subjects are strongly associated with specific taxonomic and functional signatures. Other subjects show mixed contributions, reflecting more complex microbiome organization. Importantly, some subjects belonging to the same taxonomic signature can differ substantially at the functional level, highlighting limitations of analyses based solely on taxonomy.

**Figure 3.**
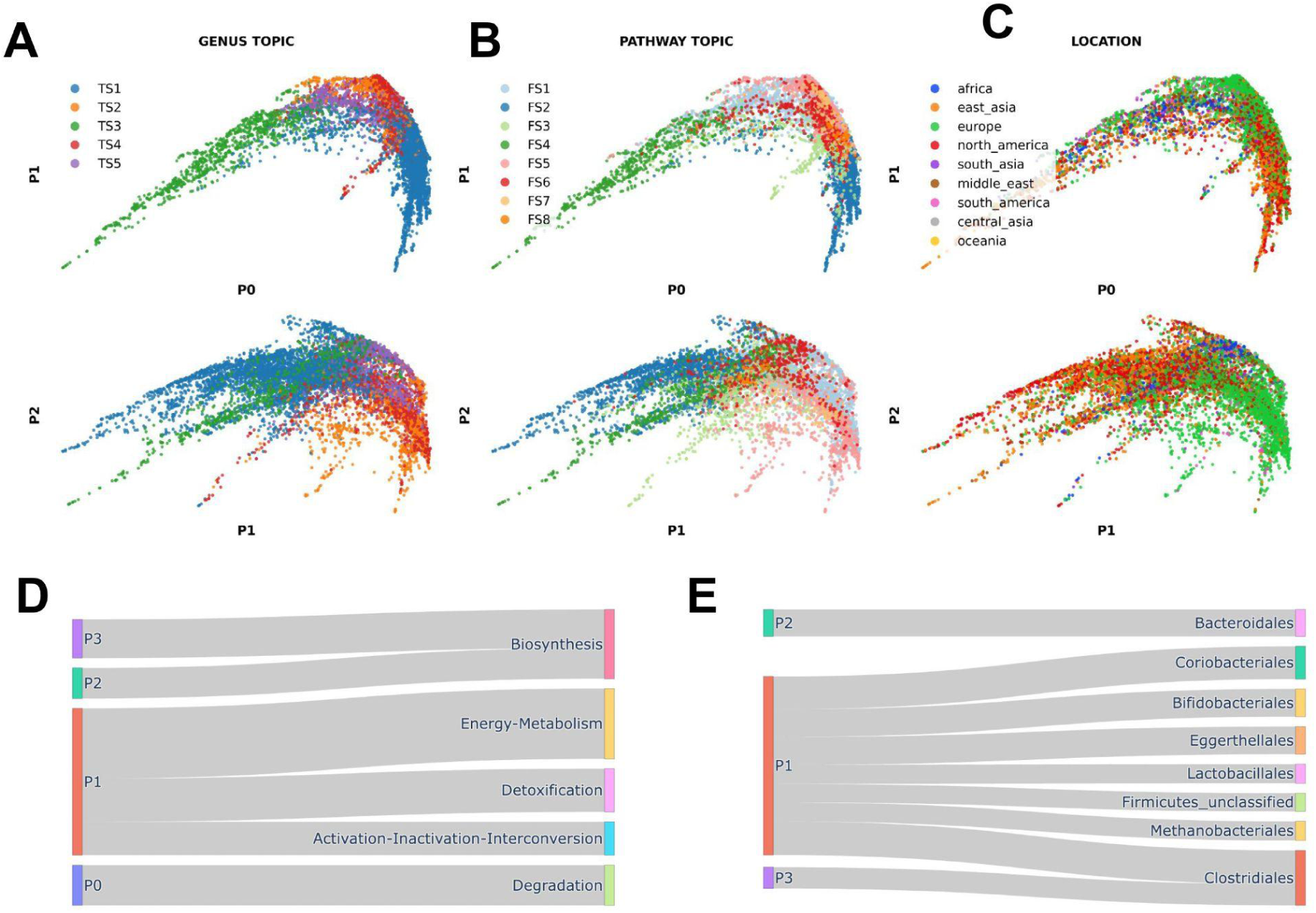
Latent representation learned by the GUT-FORMer model. A. Dimensionality reduction using the PHATE method. Each sample represents an embedded subject. Samples are colored by a dominant taxonomic signature. B. Samples colored by functional signatures. C. Samples colored by geographic location. D. Sankey plot representing pearson correlation between 5 PHATE components and pathways on aggregated to subcategory level. Only correlations above 0.3 are shown. E. Sankey plot representing pearson correlation between 5 PHATE components and taxonomy aggregated to family level. Only correlations above 0.3 are shown.

PHATE coordinates were driven primarily by biological signal, as the NMF-derived taxonomic and functional signatures explained 69.6% and 61.0% of variance in the primary PHATE coordinate (P0), respectively; study of origin accounted for only 4.6% and 7.3% of residual variance after adjusting for each signature (partial R² = 0.046 and 0.073), demonstrating that the latent representation learned by GUT-FORMer is not substantially confounded by batch effects and captures biological signals rather than technical variation (supplementary Figure 3A).

Geographic location is a major factor influencing gut microbiome composition, consistent with previous observations that microbiomes vary across populations (Andreu-Sánchez et al. 2025; Gaulke and Sharpton 2018). This has important implications for microbiome-based diagnostics and interventions, as therapeutic strategies may need to be tailored to specific populations. Influence of geographic location on gut microbiome composition is reflected in the latent representation (Fig. 3C). Without stratifying by geographic location, separation between healthy and diseased subjects using only two dimensions is less distinct which might suggest higher complexity of the connection between health and gut microbiome.

Finally, the latent representation shows that subjects within the same signature are not identical—they can still differ in their dominant species or pathways, reflecting the heterogeneity of the gut microbiome (Supplementary Fig. 3C–F).

#### Latent interpretation

Pearson correlation analysis between each latent dimension and specific features revealed that individual latent dimensions are associated with particular bacterial taxa and functional pathways. To simplify visualization, taxonomic data were collapsed to the family level and functional data to the functional subcategory level. A large proportion of latent dimensions correlate with biosynthesis-related pathways, while another group is associated with energy metabolism. Interestingly, latent dimension 1 is connected to the largest number of superpathways (Fig. 3D).

PHATE components correlate with separate subcategories of metabolic processes. P0 correlates uniquely with pathways belonging to degradation subcategories. P2 and P3 correlate with Biosynthesis and P1 correlates with multiple subcategories (Fig. 3D). On the other hand, when analyzing correlation between PHATE components and bacterial families, P1 correlated with multiple bacterial families, while P2 correlates strongly only with Bacteroidales and P3 with Clostridia. Detailed correlations between latent dimensions and individual species and pathways can be found in the Supplementary Material (Supplementary Fig. 3F and G).

Using the FACTM method (Łazęcka and Szczurek 2025), we show that specific latent dimensions are associated with disease states. Some diseases are strongly linked to individual latent dimensions, suggesting distinct microbiome alterations. Other diseases show associations across multiple latent dimensions, reflecting more complex or heterogeneous microbiome changes. Interestingly, several dimensions correlate with healthy controls, suggesting that individuals can still be considered healthy despite substantial variation in gut microbiome composition (Supplementary Fig. 3H).

Taken together, these results indicate that the GUT-FORMer latent space captures biologically meaningful structure in the data, linking microbial composition and functional potential with host-associated factors.

### GUT-FORMER REPRESENTATIONS CAN BE USED FOR DISEASE CLASSIFICATION TASKS

Next, we sought to assess whether the learned deep representation of the human gut microbiome could improve performance in disease classification and regression tasks. To this end, we evaluated binary classification between healthy and diseased subjects, multiclass classification, and regression tasks predicting BMI and age in both healthy and diseased individuals, including adults and children.

#### Binary disease classification

To evaluate the utility of the learned representation for classifying disease-associated microbiome samples, we evaluated classification performance between samples from a given disease and healthy controls. A total of 25 diseases were included, each analyzed separately using a random forest classifier within a 5-fold cross-validation framework, with grid search–based hyperparameter tuning. Only diseases with a minimum of 15 samples were considered (see Supplementary Figure 4 for the complete overview of diseases). GUT-FORMer successfully classifies multiple diseases using the learned latent representation (Fig. 4A). The model achieves a high AUC cross diseases, demonstrating discriminative performance. Classification precision and recall are also high across diseases (Fig. 4B), indicating balanced and reliable predictions.

**Figure 4.**
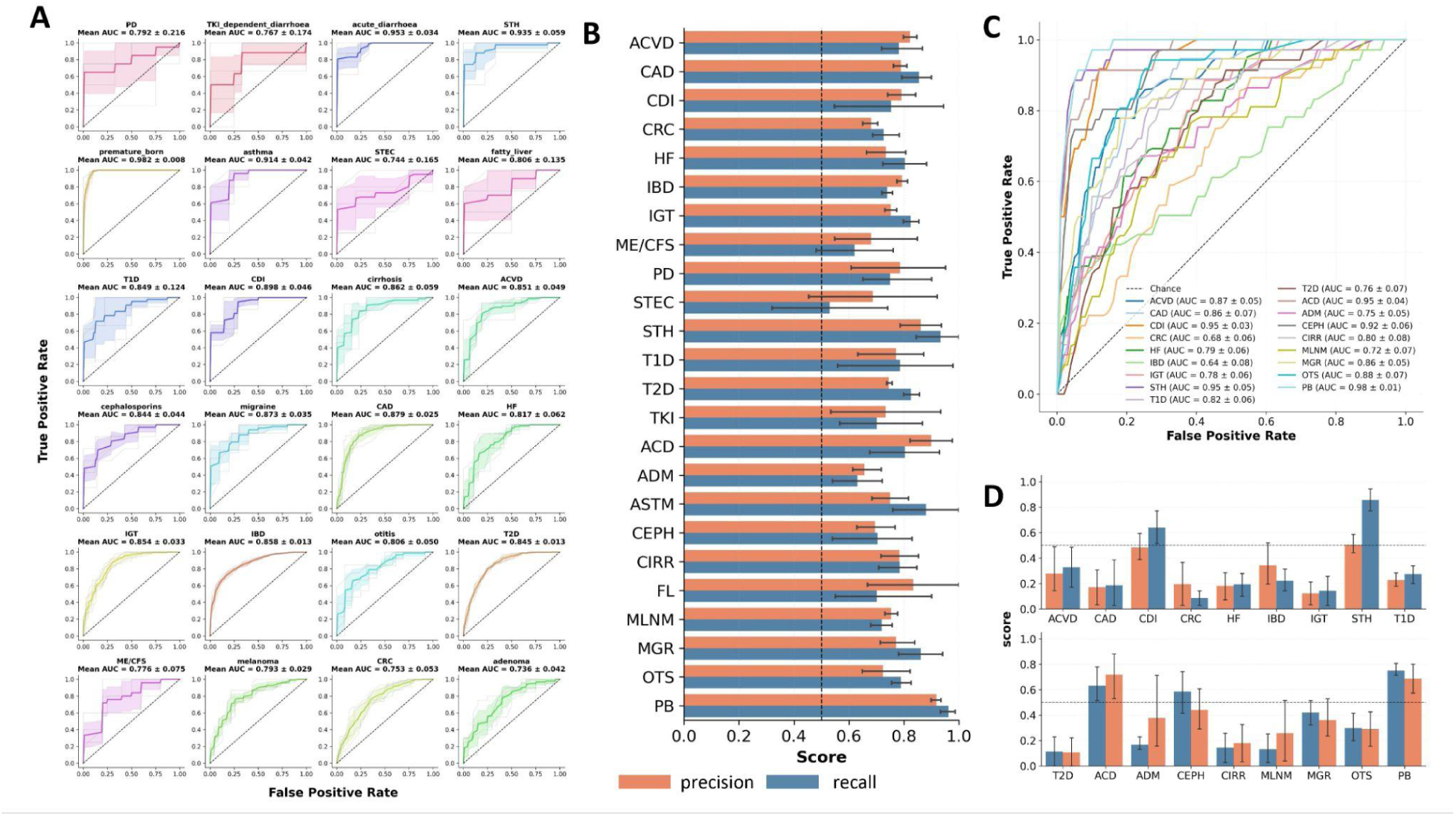
GUT-FORMer representation can be used to predict health states of the human gut microbiome. A. ROC-AUC curves showing results of k-fold binary classification between samples belonging to specific disease and any other disease. B. Precision and recall results of the binary classification between samples belonging to specific disease and any other disease. C. . ROC-AUC curves showing results of binary classification between samples belonging to specific disease and any healthy subjects. D. Precision and recall results of the binary classification between samples belonging to specific disease and any healthy sample. TKI – TKI-dependent diarrhoea; ACD – acute diarrhoea; ADM – adenoma; ASTM – asthma; CEPH – cephalosporin exposure; CIRR – cirrhosis; FL – fatty liver; MLNM – melanoma; MGR – migraine; OTS – otitis; PB – premature birth; CSH – carcinoma surgery history; ACVD – atherosclerotic cardiovascular disease; BD – Behçet’s disease; CAD – coronary artery disease; CDI – Clostridioides difficile infection; CRC – colorectal cancer; HF – heart failure; IBD – inflammatory bowel disease; IGT – impaired glucose tolerance; ME/CFS – myalgic encephalomyelitis/chronic fatigue syndrome; PD – Parkinson’s disease; STEC – Shiga toxin–producing Escherichia coli; STH – soil-transmitted helminths; T1D – type 1 diabetes; T2D – type 2 diabetes.

#### Multiclass classification

A total of 13 diseases were included in the multiclass classification analysis. Classification was performed using a random forest model with 5-fold cross-validation and hyperparameter tuning via grid search. Only diseases with at least 50 samples were considered.

GUT-FORMer achieved a high overall AUC, indicating strong performance in distinguishing among multiple disease states simultaneously (Fig. 4C). Precision and recall exhibited greater variability compared to the binary classification setting, reflecting the increased complexity of the multiclass task (Fig. 4D). Nevertheless, overall performance remained high, suggesting that the learned latent representation effectively captures disease-specific microbiome structure.

### GUT-FORMER REPRESENTATIONS CAN BE USED FOR REGRESSION TASKS

#### Age prediction

Age was predicted using the latent representation as input and a RandomForestRegressor with a five-fold cross-validation approach and hyperparameter tuning via grid search. Age distribution in samples can be found in Supplementary Fig 5A.

GUT-FORMer accurately predicts host age in both healthy and diseased adults, demonstrating that microbiome representations capture age-associated biological variation (Fig. 5A). Age prediction performance is slightly higher in diseased individuals compared to healthy controls (Pearson correlation coefficient *r* = 0.37 and R² = 0.13 for healthy individuals, and *r* = 0.49 and R² = 0.22 for diseased individuals), suggesting that disease-associated microbiome changes may follow more consistent or accelerated trajectories.

**Figure 5.**
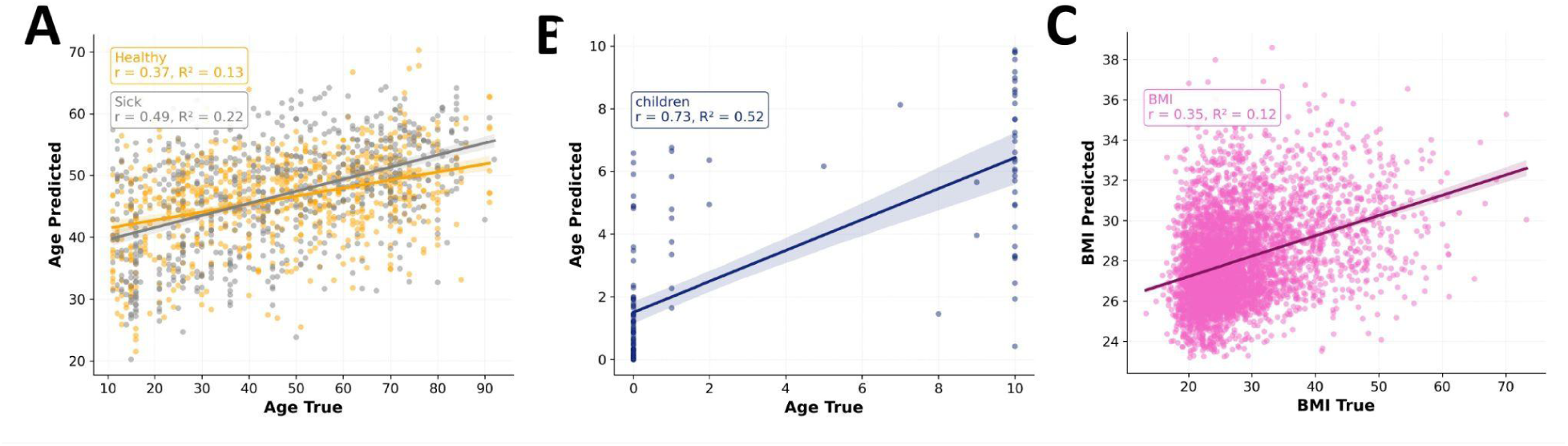
GUT-FORMer representation is useful in regression tasks such as age and BMI prediction. A. Age prediction for adult subjects separately for healthy and diseased subjects. Scatterplot representing true age and age predicted by RandomForest regressor trained on GUT-FORMer representation. B. Age prediction for healthy children C. BMI prediction for adult subjects. Scatterplot representing true BMI and BMI predicted by RandomForest regressor trained on GUT-FORMer representation.

The model also predicts age in children with strong performance (Fig. 5B), achieving a Pearson correlation coefficient of *r* = 0.73 and a coefficient of determination of R² = 0.52.

#### BMI prediction

BMI was predicted using the latent representation as input and a RandomForestRegressor with a five-fold cross-validation approach and hyperparameter tuning via grid search.The model predicts BMI with moderate accuracy, achieving a Pearson correlation coefficient of *r* = 0.35 and a coefficient of determination of R² = 0.12 (Fig. 5C). BMI distribution in samples can be found in Supplementary Fig 5B.

Our model outperforms previously reported microbiome-based age prediction approaches. Even though we didn’t find a paper that separately predicts age for sick and healthy individuals, Huang et al. (Huang et al. 2020) reported an R² of 0.16, while Zahavi et al. (Zahavi et al. 2025) achieved an R² of 0.14. In comparison, GUT-FORMer reaches higher predictive performance, particularly in diseased individuals and in the pediatric cohort, highlighting the advantage of learned latent representations for capturing age-associated variation. A similar trend was observed for BMI prediction, where the latent representation learned by GUT-FORMer yielded an R² of 0.12, compared to 0.16 reported by Zahavi et al. (Zahavi et al. 2025). Despite the slightly lower explained variance, this result remains comparable and supports the utility of the learned representation in capturing relevant host–microbiome associations.

These results demonstrate that the GUT-FORMer latent representation captures meaningful host-associated biological signals, including age and BMI.

#### GUT-FORMer OUTPERFORMS OTHER MODELS IN DOWNSTREAM TASKS

To benchmark the GUT-FORMer latent representation of the human gut microbiome and its utility in classification and regression tasks, we compared it against several established methods: DeepMicro (Oh and Zhang 2020), GMHI (Gupta et al. 2020), Dysbiosis Index (Zielińska et al. 2025) and taxonomic NMF (Frioux et al. 2023) The Gut Microbiome Health Index (GMHI) is a metric designed to quantify overall gut health based on the relative abundance of health- and disease-associated microbial species. The Dysbiosis Index (Q2PD) is a function-centric metric that quantifies the degree of microbiome imbalance by evaluating species-level functional potential and interspecies interactions. DeepMicro is a deep learning framework that learns low-dimensional representations of microbiome profiles to improve disease classification and other predictive tasks. We also included functional signatures to this benchmark. Additionally, we performed model ablations to further evaluate performance, including versions of GUT-FORMer trained solely on taxonomic or functional data.

We first evaluated the learned embeddings in a binary classification task that aims to separate healthy and diseased subjects. Interestingly, GUT-FORMer trained solely on functional information achieved the best performance (Fig. 6A; Supplementary Fig. 6A,B). Precision–recall analysis further showed that the functional-only model performed comparably to the model trained on both taxonomic and functional inputs (Fig. 6B). Performance was evaluated for individual diseases, can be found in the Supplementary Fig. 6B.

**Figure 6.**
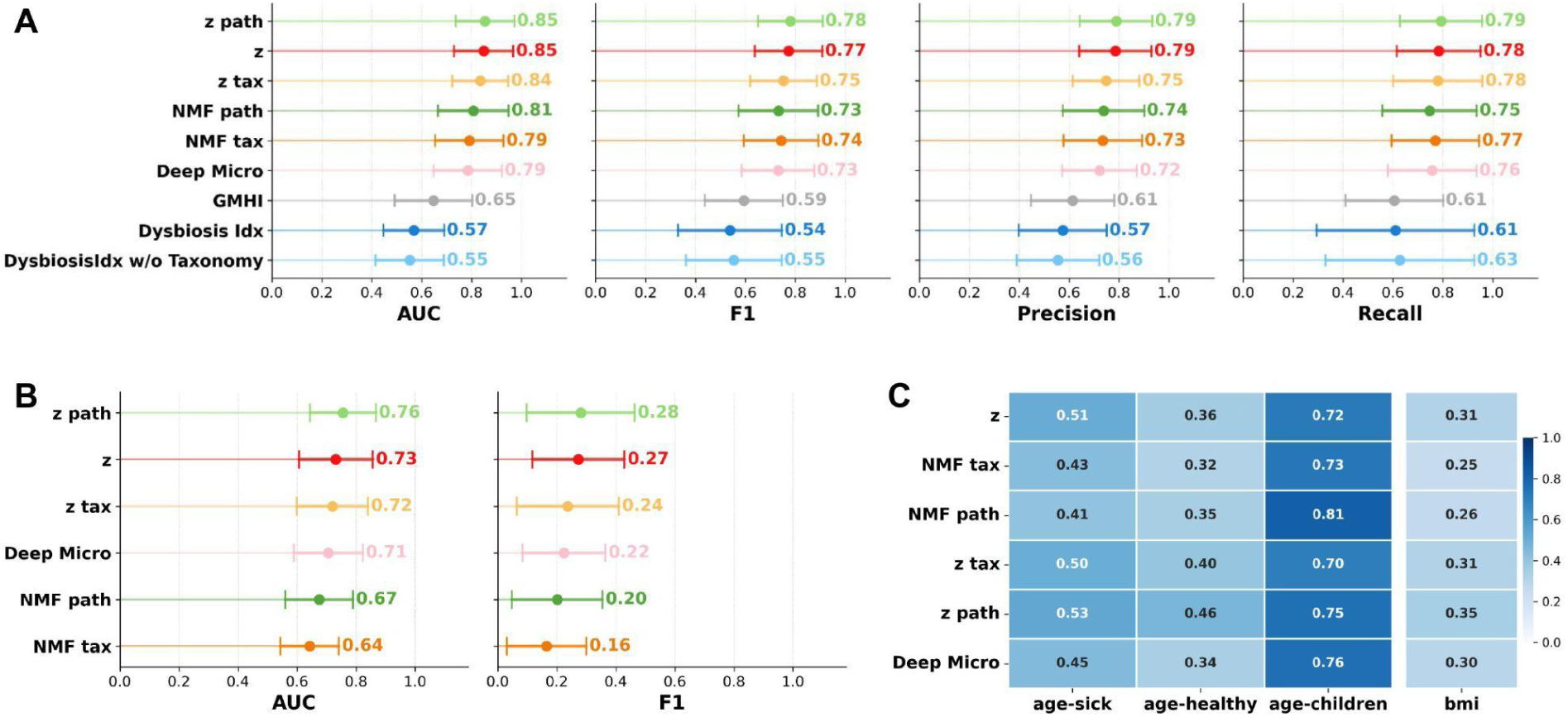
Benchmarking GUT-FORMer results for classification and regression tasks using 5-fold cross-validation and grid search. A. Binary classification of diseased versus healthy individuals using a random forest model. B. Multiclass classification performance. C. Regression performance for predicting age in diseased adults, healthy adults, and healthy children, as well as BMI prediction.

We next evaluated GUT-FORMer in a multiclass classification setting. For this analysis, GMHI and functional indices were excluded, as they are not applicable in this context. Consistent with the binary classification results, GUT-FORMer outperformed all benchmarked methods. Notably, the functional-only ablation achieved performance comparable to the full model across diseases (Fig. 6B). However, compared to the binary setting, the F1 score was lower, reflecting the increased difficulty of the multiclass task, where the model must discriminate among multiple disease states simultaneously.

We further evaluated GUT-FORMer in regression tasks, including age prediction in healthy and diseased adults, age prediction in healthy children, and BMI prediction. In all cases, GUT-FORMer outperformed benchmark models, with the full model trained on both taxonomy and pathways showing superior performance compared with the functional-only ablation (Fig. 6C; Supplementary Fig. 6D–G).

Together, these results indicate that the representations learned by GUT-FORMer capture biologically meaningful structure within the human gut microbiome and outperform existing approaches in distinguishing between healthy and diseased states.

GUT-FORMer representation trained exclusively on functional profiles often matched or even exceeded the performance of the mode integrating both taxonomic and pathway-level information, highlighting the strong predictive signal contained in functional data. We hypothesize that this observation is related to the lower sparsity of functional profiles in the human gut microbiome. In contrast to taxonomic abundance matrices, functional profiles are less sparse, which may reflect the substantial proportion of the gut microbiome that remains taxonomically uncharacterized or unannotated. This reduced sparsity may provide a more stable and informative representation for downstream learning tasks. We further hypothesize that with increasing sample sizes and improved taxonomic resolution, the relative performance advantage of functional data may diminish. Nevertheless, we believe that incorporating taxonomic information remains valuable, as it enhances interpretability and yields the best overall performance in regression-based tasks.

## DISCUSSION

Understanding the human gut microbiome remains challenging due to its compositional nature, high sparsity, and substantial inter-individual variability (Kumar et al. 2024). Here, we introduce two complementary approaches to better characterize population structure in gut microbiome data: functional signatures of the human gut microbiome and GUT-FORMer, a transformer-based autoencoder that integrates taxonomic composition and functional potential to learn microbiome representations and predict host phenotypes.

We show that functional signatures of the gut microbiome reveal patterns that are not always apparent at the taxonomic level. These signatures capture ensembles of metabolic pathways rather than individual taxa, reflecting coordinated microbial activity. While many functional signatures align with specific taxonomic patterns, several are largely independent of taxonomy, indicating that distinct microbial communities can achieve similar functional states. This finding reinforces the complementary nature of taxonomic and functional perspectives and highlights the limitations of taxonomy-only approaches for understanding microbiome-mediated health effects (Zielińska et al. 2025). The developed approach may help address ongoing questions regarding functional redundancy in microbial systems, which in turn raises fundamental issues about the mechanisms enabling species coexistence and the role of diversity in community functioning (Louca et al. 2018). It also relates to the broader question of what more accurately defines microbial communities: “who is there” versus “what they can do” (Xu et al. 2014). Resolving this debate is particularly important in the context of live biotherapeutic development and administration, where functional stability and metabolic output may be more directly relevant than taxonomic composition alone.

Understanding these relationships may therefore inform the rational design and selection of microbial consortia with desired functional properties, improving the precision and efficacy of microbiome-based therapeutic interventions (Min et al. 2025; Konar 2026; Brevi and Zarrinpar 2023).

Building on these insights, GUT-FORMer provides a unified latent representation of the microbiome by jointly modeling taxonomic and functional data with a transformer-based autoencoder architecture. The learned embeddings capture biologically meaningful structure, reflect previously defined functional and taxonomic signatures (Frioux et al. 2023), and reveal additional patterns beyond classical NMF-derived signatures. The latent space distinguishes healthy from diseased subjects, captures population-specific variability, and identifies latent dimensions associated with particular microbial taxa, pathways, and disease states.

Importantly, GUT-FORMer supports downstream predictive tasks. It achieves high performance in binary and multiclass disease classification, as well as regression tasks including host age and BMI, outperforming existing microbiome indices and deep learning methods. These results demonstrate that integrating taxonomy and function produces a more informative and generalizable representation than relying on either modality alone. Functional-only models capture strong predictive signals, particularly for disease classification, but including taxonomic information improves interpretability and enables linking functional potential to specific microbial taxa—a critical requirement for microbiome-targeted interventions.

Our findings have direct implications for precision medicine. Functional signatures and latent representations allow classification of individuals into microbiome-based subgroups, which can inform personalized probiotic strategies, dietary interventions, or therapeutic response predictions. The model also highlights population-specific variation, emphasizing that microbiome-based diagnostics or interventions may require adjustment for geographic or demographic context.

Despite these advances, several limitations remain. Functional profiling captures only a subset of the microbiome’s metabolic potential due to incomplete annotation of microbial genes and proteins (Maranga et al. 2023). Predicted functional profiles represent potential rather than realized activity, and integrating metatranscriptomic or metabolomic data could further improve functional resolution. Microbiome composition is shaped by environmental and host factors, and although our dataset is large and diverse, it may not fully capture global variation (Gacesa et al. 2022). Finally, while GUT-FORMer learns biologically meaningful representations, deep learning models remain less interpretable than classical methods, and additional work is needed to dissect which specific features drive predictions(Rudin 2019).

In summary, this study demonstrates that joint modeling of taxonomic and functional data provides a powerful framework for microbiome analysis, uncovering biologically relevant patterns, improving predictive performance, and supporting translational applications. GUT-FORMer establishes a generalizable platform for future studies of microbiome-mediated health, disease, and therapeutic response, moving the field closer to precision microbiome medicine.

## ACKNOWLEDGEMENTS

This research was funded by the project of the Minister of Science and Higher Education “Support for the activity of Centers of Excellence established in Poland under Horizon 2020” on the basis of the contract number MEiN/2023/DIR/3796.

The authors thank prof. Anna Karnkowska (Institute of Evolutionary Biology, University of Warsaw, Poland) for her Master thesis supervision of A.R.

Merck Healthcare KGaA provides funding for the Szczurek-lab research group (M.M and E.S.).

## Data availability

The code is available on GitHub at: https://github.com/Tomasz-Lab/gut-former

## Author contributions

Z.K. and T.K. conceived the study; T.K. and E.S. supervised the research; Z.K., T.K., A.R., M.M., E.S. analysed the data; Z.K. implemented the original version of GUT-FORMer; A.R. performed and analyzed enterosignatures; W.N. implemented the production code-base; E.S., M.M. consulted on the choice of machine learning models and statistical approaches; Z.K. and T.K. wrote the paper. All authors revised and approved the paper.

## METHODS

All code was implemented in Python (version 3.9.19). <u>MOTIVATION AND MODEL OVERVIEW</u>

### Datasets

Publicly available datasets from *CuratedMetagenomicData (Pasolli et al. 2017)* were used in this study. Both taxonomic profiles (provided as .relative_abundance) and functional profiles (provided as .pathway_abundance) were analysed. The functional data are originally stratified by bacterial species; therefore, for each pathway, abundances were aggregated across all species and summed to obtain total pathway abundance. A complete list of the datasets included in the analysis is provided in Supplementary Figure 1. Only stool samples were considered. Samples were filtered to retain only disease groups with more than three subjects. The dataset was then split into training and validation sets (90/10), using stratification by age category within each disease group where possible; otherwise, random splitting was applied.

### Data preprocessing

Taxonomic and functional profiles were preprocessed by filtering low-prevalence features, retaining only those present in at least 10% of samples. The data were subsequently normalised to relative abundances per sample and multiplied by 100. Functional profiles were multiplied by a factor of 10.

## FUNCTIONAL SIGNATURES OF THE HUMAN GUT MICROBIOME

### Non-Negative Matrix Factorization

NMF is a matrix factorization technique that decomposes a non-negative input matrix into two lower-dimensional non-negative matrices (W and H), capturing latent components that represent additive patterns in the data. The implementation was based on sklearn.decomposition.NMF from Scikit-Learn v1.8.0.

### NMF parameters

NMF was performed with class sklearn.decomposition.NMF from Scikit-Learn v1.8.0 using the multiplicative update solver, Kullback-Leibler divergence as a beta-loss function, random initialization and a maximum iterations of 2000. The best model from 100 runs was chosen based on the highest explained variance and cosine similarity. Parameter of regularization set to 0 (no regularization).

### Defining the number of k

In order to determine the number of signatures, nine-fold bicross-validation was performed as described in Owen and Perry, Kanagal and Sindhwani, and Eng et al.. All runs were performed for each number of clusters k, ranging from 2 to 15. The quality of decomposition for the validation was calculated using explained variance and cosine similarity. The optimal number of signatures was chosen by assessing the gain in cosine similarity (mean for all runs) when increasing the number of signatures: the point after which the curve stops rising steeply and starts to flatten out was selected as the optimal number of signatures (KneeLocator from kneed package).

### Verification of stability

To verify the stability of the NMF algorithm, the resulting H and W matrices from 50 runs were tested to determine whether the composition of signatures changes significantly for each sample/function across runs. First, pairwise euclidean distances were calculated across all runs for a given sample/function. Then, each sample/function composition of topics was shuffled and distances were recalculated. Based on the resulting mean p-value for all samples/functions (for number of permutations is 0.0099), we can reject the null hypothesis that topic contributions are different across runs for a single sample/function.

### Dominant features in each signature

To analyze the contribution of each pathway to functional signatures, the H matrix was first transformed into relative abundance. Feature contributions within each signature were then assessed and visualized. The five most abundant features per signature were selected for plotting. To analyze signature abundance per subject, the W matrix was transformed into relative abundance. The dominant signature for each subject was assigned based on the highest abundance.

### Signatures visualization

PCoA was computed on the functional matrix using Aitchison distance with a pseudocount of 1e-3. Both the Aitchison distance computation and the PCoA embedding were performed using scikit-bio (0.6.2). Dominant functional signatures were visualized on the resulting ordination plot.

### Correlation with bacterial genera

The taxonomy matrix was collapsed to the genus level by aggregating species-level abundances. Pearson correlation was then computed between each genus and each functional signature using SciPy (1.13.1).

### Taxonomy-based enterosignatures

Taxonomy-based enterosignatures were calculated based on Frioux et al. (Frioux et al. 2023) Species-level taxonomy matrices were aggregated to the genus level and transformed into relative abundance. Five NMF components were selected. Both W and H matrices were transformed into relative abundance. The dominant signature per subject was assigned based on abundance, and the dominant genus per signature was identified based on feature contribution. The distribution of dominant signatures and the abundance of each taxonomic feature across diseases are shown in Supplementary Fig. 2C.

### Correlation between taxonomic and functional signatures

Pearson correlation between taxonomic and functional signatures was computed using scipy.stats.pearsonr, which returns both the correlation coefficient and the associated p-value.

## GUT-FORMER LATENT SPACE CAPTURES BIOLOGICALLY RELEVANT STRUCTURE IN THE HUMAN GUT MICROBIOME

### GUT-FORMer training

A deep learning model based on the Transformer architecture was developed to jointly model taxonomic and functional microbiome profiles. The model was implemented and trained using PyTorch (2.4.1). Separate embedding layers were used to encode bacterial taxa and metabolic pathways. These embeddings were combined with abundance information and processed using Transformer encoder layers to capture complex relationships across features. The model learns a shared latent representation, which is subsequently decoded to reconstruct both taxonomic and functional profiles. The model was trained using paired taxonomic and pathway abundance data, split into training and test sets (90/10). Input features were scaled prior to training. Optimization was performed using the Adam optimizer with a weight decay of 0.001.

The training objective combined multiple components: (i) reconstruction loss (mean squared error) for both taxonomic and functional outputs, (ii) a Log Euclidean distance preservation loss to maintain pairwise relationships between samples, and (iii) a variance regularization term to preserve global data structure. Model hyperparameters, including embedding dimension, latent dimensionality, learning rate, and batch size, were selected using grid search with the objective of minimizing the overall loss function. Model performance was evaluated using reconstruction error.

Training was conducted for up to 1000 epochs with early stopping based on validation loss to prevent overfitting. The model with the best validation performance was retained, and performance metrics were recorded throughout training.

Final training was performed using the following parameters: batch size of 16, learning rate of 1.893292917167e-4, embedding dimension of 128, latent dimension of 64, one Transformer encoder layer, and two attention heads. The loss function was weighted as follows: reconstruction weight = 1, distance weight = 0.1, and variance weight = 0.1.

### Ablations

The GUT-FORMer model was trained using the same parameters, with the modification of using only a single modality (either taxonomy or pathways) as input.

### Reconstruction error calculation

Reconstruction performance was evaluated using mean squared error (MSE) and Procrustes disparity, computed separately for taxonomic and functional matrices. Procrustes disparity was computed using SciPy (1.13.1).

MSE is defined as the average of the squared differences between the original and reconstructed values across all samples and features.

Procrustes (Eguizabal et al. 2019) analysis is a shape comparison method that assesses the similarity between two configurations (here, sample ordinations) after optimal scaling, translation, and rotation. The Procrustes disparity is defined as the sum of squared distances between corresponding points in the aligned configurations, providing a measure of dissimilarity between the original and reconstructed sample structures.

### PHATE dimensionality reduction

The 64-dimensional GUT-FORMer latent representation was reduced for visualization using PHATE with default parameters, implemented in phate (2.0.0). PHATE is a nonlinear dimensionality reduction method that preserves both local and global structure in high-dimensional data by modeling diffusion-based transitions between data points. Three components were visualized using scatterplots (matplotlib 3.9.2).

Dominant taxonomic and functional signatures were projected onto the PHATE embedding. Additional metadata were visualized, including dominant species and pathways per subject, country, study, and health status.

Batch effects were assessed in the PHATE latent space using the first component (P0). For each taxonomic and functional signature, ordinary least squares models were fitted using statsmodels (0.14.3), comparing (i) models with the signature alone and (ii) models including both the signature and study (categorical, sum contrasts). The unique contribution of study was quantified as the increase in explained variance in the full model relative to the condition-only model, scaled by the remaining unexplained variance, representing variance attributable to study beyond the biological signal. Supplementary Fig. 3A–E.

### Latent dimension interpretation

Pearson correlation analysis between each latent dimension and input features was performed using SciPy (1.13.1), showing that individual dimensions are associated with specific bacterial taxa and functional pathways. For visualization, taxonomic data were aggregated to the family level and functional data to the subcategory level. Correlations at the species and pathway level are provided in Supplementary Fig. 3F–G.

Association between latent dimensions and diseases was assessed using the FACTM method (Łazęcka and Szczurek 2025), a multivariate statistical approach that evaluates the relationship between continuous latent variables and categorical outcomes while accounting for covariance structure across features. Latent factors were aligned to external variables to improve interpretability using a rotation-based approach. A dependence matrix between latent factors and external variables was computed using pairwise Pearson correlations, excluding missing values per variable. This matrix was decomposed using singular value decomposition (SVD), implemented in NumPy (2.0.2), to derive an orthogonal rotation, which was subsequently applied to the latent factors. To ensure compatibility between latent and external representations, the external variable matrix was padded with Gaussian noise where required. Alignment was assessed by comparing correlation structures before and after rotation.

Associations between rotated latent factors and clinical outcomes were evaluated using the Kruskal–Wallis test, implemented in SciPy (1.13.1). Multiple hypothesis testing was controlled using false discovery rate correction via SciPy (1.13.1).

## GUT-FORMER REPRESENTATIONS CAN BE USED FOR DISEASE CLASSIFICATION TASKS

### Binary classification

Validation samples were used for this analysis. Diseases with fewer than 15 samples were excluded. Input data consisted of GUT-FORMer latent representations from validation samples. For each disease, a random forest classifier was trained using 5-fold cross-validation with hyperparameter tuning via grid search, implemented in scikit-learn (1.5.2). In each fold, diseased samples were compared against healthy controls. Predicted disease labels are shown in Supplementary Fig. 4.

Model performance was evaluated using area under the ROC curve (AUC), F1 score, precision, and recall, computed using scikit-learn (1.5.2). AUC (area under the receiver operating characteristic curve) is defined as the probability that the model ranks a randomly chosen positive sample higher than a randomly chosen negative sample. Precision is defined as the proportion of true positive predictions among all positive predictions. Recall (sensitivity) is defined as the proportion of true positive predictions among all actual positive samples. F1 score is defined as the harmonic mean of precision and recall.

### Multiclass classification

Diseases represented by fewer than 50 samples were excluded from the analysis. For each remaining disease, a random forest classifier was trained on GUT-FORMer latent representations using scikit-learn (1.5.2). Model performance was evaluated using 5-fold cross-validation, with hyperparameters optimized via grid search. Predictive performance was assessed using area under the ROC curve (AUC), F1 score, precision, and recall, computed with scikit-learn (1.5.2).

## GUT-FORMER REPRESENTATIONS CAN BE USED FOR REGRESSION TASKS

### Age prediction in adults

Adults were defined as subjects older than 10 years. GUT-FORMer representations were used as input to a random forest regressor implemented in scikit-learn (1.5.2). Models were trained using 5-fold cross-validation. Performance was evaluated using Pearson correlation and coefficient of determination (R²), computed with scikit-learn (1.5.2). R² is defined as the proportion of variance in the observed data explained by the model. Models were fitted separately for healthy and diseased subjects.

### Age prediction in children

Children were defined as subjects younger than 10 years. The same modeling procedure was applied.

### BMI prediction

Children (age <20) were excluded. GUT-FORMer representations were used to predict BMI using a random forest regressor implemented in scikit-learn (1.5.2), with 5-fold cross-validation. Performance was evaluated using Pearson correlation and coefficient of determination (R²), computed with scikit-learn (1.5.2).

## GUT-FORMer OUTPERFORMS OTHER MODELS IN DOWNSTREAM TASKS

### Analyzed models

The following models were evaluated: GMHI, Functional Dysbiosis Index, DeepMicro, NMF-based representations (taxonomy and function), and GUT-FORMer representations trained on taxonomy only and function only.

Non-negative matrix factorisation (NMF) was trained on the same training dataset as GUT-FORMer, implemented in scikit-learn. The fitted model was subsequently used to transform the validation data into the learned latent space.

GMHI (Gut Microbiome Health Index) is a composite index derived from taxonomic profiles that quantifies microbiome health based on the relative abundance of health-associated and disease-associated microbial species (Gupta et al. 2020) .

Functional Dysbiosis Index is a metric based on functional pathway profiles that quantifies deviations from a healthy microbiome by comparing the relative abundance of pathways associated with health and disease states (Zielińska et al. 2025). This index was calculated using both a function-only approach and a combined taxonomy–function approach.

DeepMicro is a deep learning–based framework that learns low-dimensional representations of microbiome data (e.g., using autoencoders or variational autoencoders) for downstream tasks such as classification and regression. (Oh and Zhang 2020)

For benchmarking of the deep autoencoder (DAE), we followed the DAE architecture search space reported in the DeepMicro supplementary material. Specifically, for the species-level relative abundance profiles, we evaluated 10 DAE configurations obtained by combining five latent dimensionalities (32, 64, 128, 256, and 512) with either one or two hidden layers in both the encoder and decoder. All other network settings, including activation functions and weight initializers, were kept at their default values. The autoencoder code was modified to save both the full trained model and the encoder separately, enabling extraction of latent representations in addition to reconstruction outputs. For each tested architecture, latent embeddings and reconstructed profiles were generated for both training and validation datasets and used for comparative assessment. Based on this benchmark, the best-performing architecture was the 64–32 encoder configuration, corresponding to a latent space of size 32.

### Binary classification

Diseases with fewer than 15 samples were excluded. Input data consisted of GUT-FORMer latent representations from validation samples. For each disease, a random forest classifier was trained using 5-fold cross-validation with hyperparameter tuning via grid search, implemented in scikit-learn (1.5.2). Models were compared based on AUC, F1 score, precision, and recall averaged across folds and diseases.

### Multiclass classification

Diseases represented by fewer than 50 samples were excluded from the analysis. For each remaining disease, a random forest classifier was trained on GUT-FORMer latent representations using scikit-learn (1.5.2). GMHI and Functional index were removed from this analysis. Models were compared based on AUC, F1 score, precision, and recall averaged across folds and diseases.

### Regression

For regression tasks, GMHI and Functional Dysbiosis Index were excluded. Only ablation models, NMF-based representations, and DeepMicro were compared to GUT-FORMer. The benchmark included age prediction in sick and healthy adults and children, as well as BMI prediction in adults. Adults were defined as subjects older than 10 years, and children as subjects younger than 10 years. GUT-FORMer representations were used as input to a random forest regressor implemented in scikit-learn (1.5.2). Models were trained using 5-fold cross-validation. Performance was evaluated using Pearson correlation and coefficient of determination (R²), computed with scikit-learn (1.5.2).

