## Supplementary Information for "A universal taxonomic and functional human gut microbiome model for disease classification and phenotype discovery"

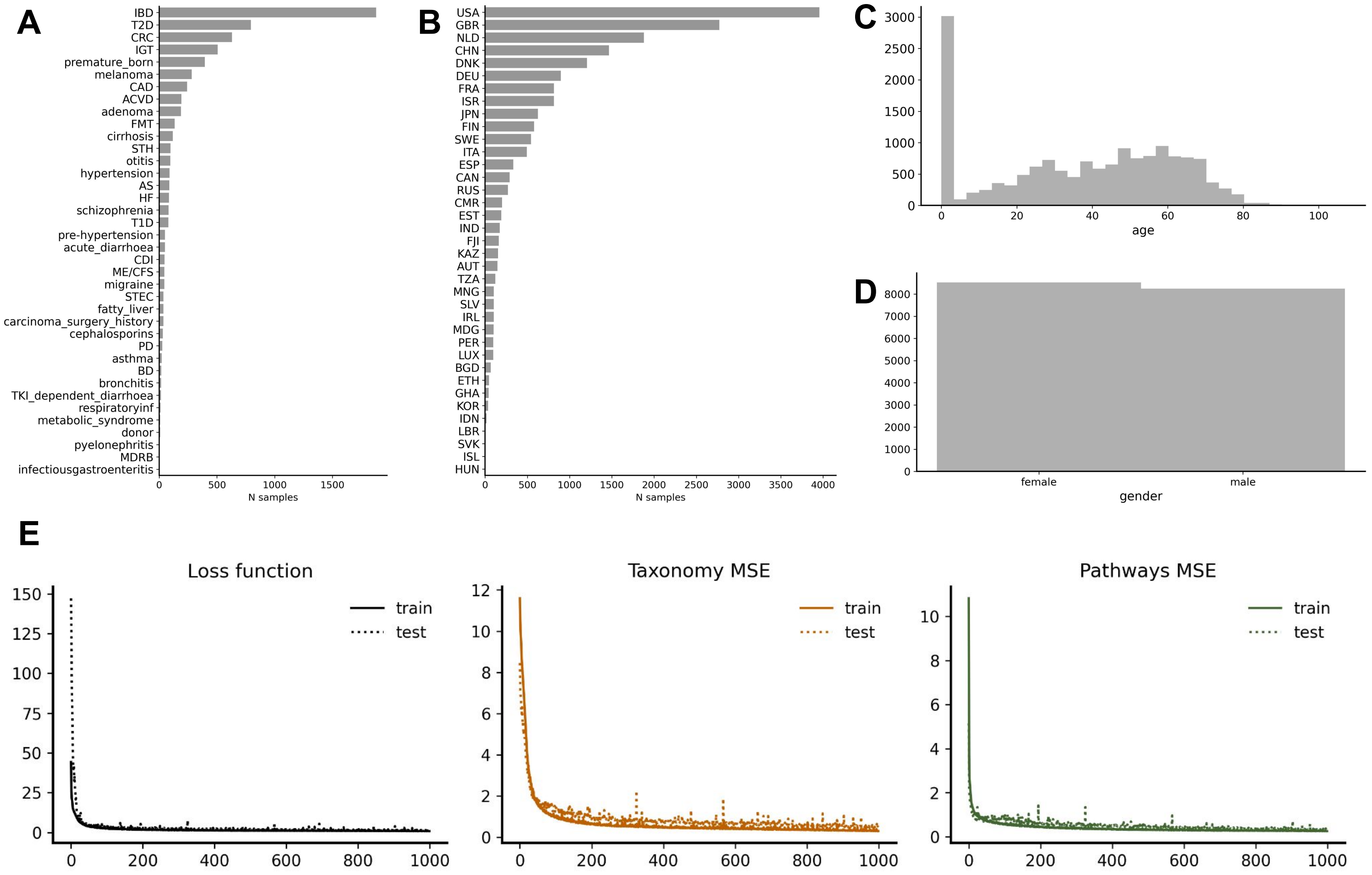


**Supplementary Figure 1. Training data overview and model performance.** A. Distribution of disease categories across samples; B. Geographic distribution of samples by country; C. Age distribution of participants; D. Gender distribution of participants; E. Training dynamics showing the loss function across epochs; F. Taxonomic reconstruction performance across epochs; G. Pathway reconstruction performance (mean squared error) across epochs.

**Supplementary Figure 2. Functional signatures overview.** A. Full feature contribution to each functional topic;

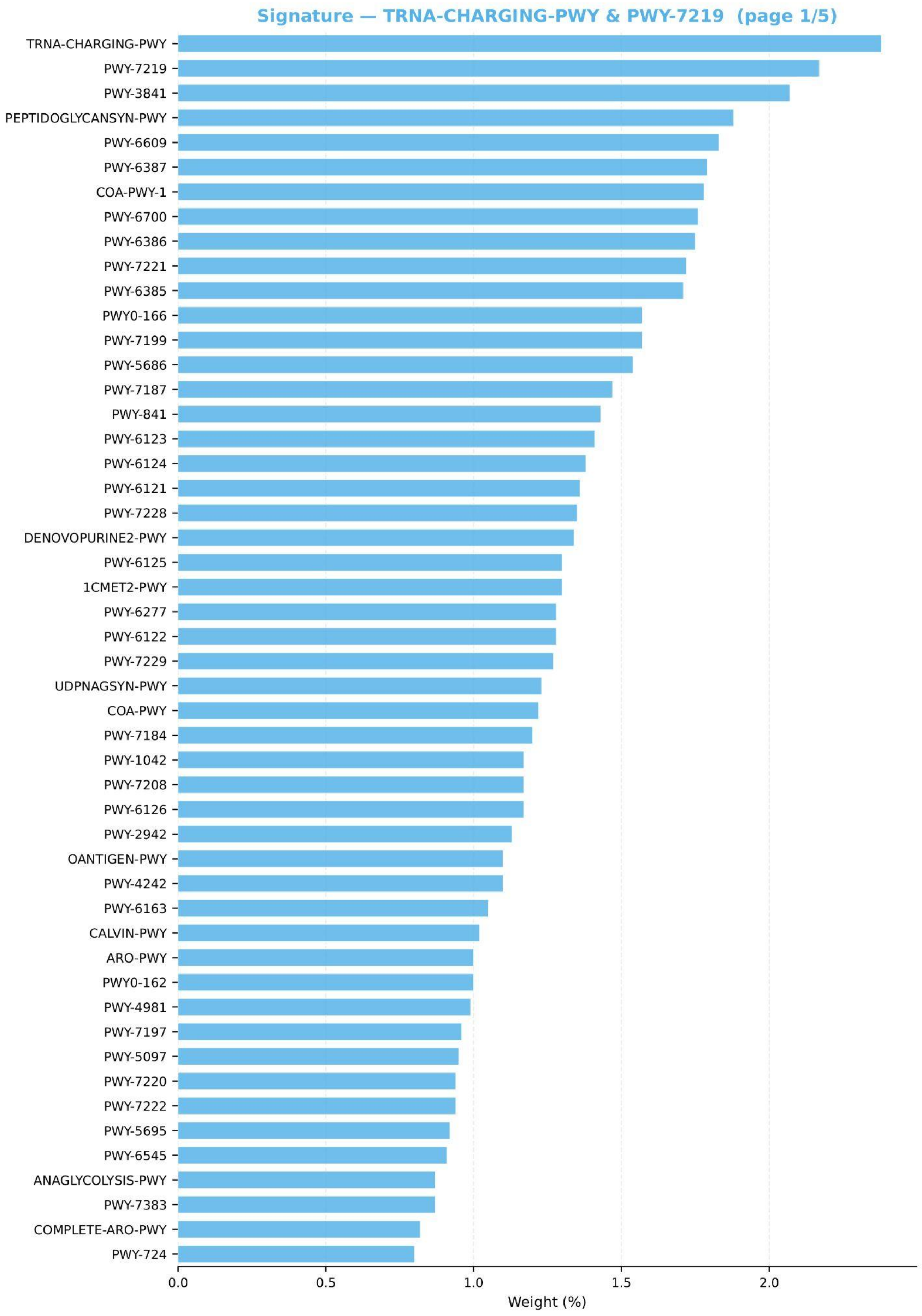

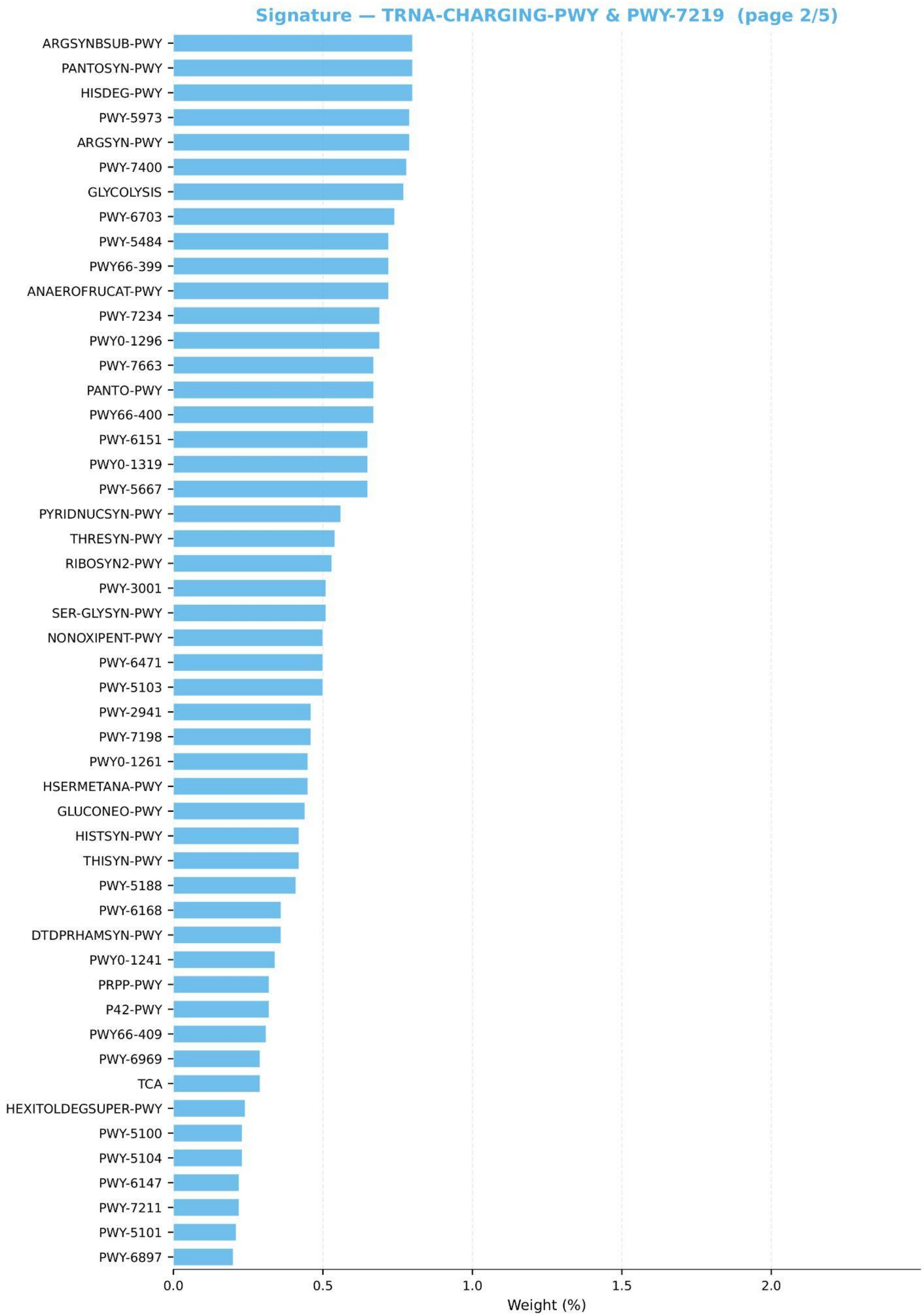
Signature — TRNA-CHARGING-PWY & PWY-7219 (page 2/5)
ARGSYNBSUB-PWY
PANTOSYN-PWY
HISDEG-PWY
PWY-5973
ARGSYN-PWY
PWY-7400
GLYCOLYSIS
PWY-6703
PWY-5484
PWY66-399
ANAEROFRUCAT-PWY
PWY-7234
PWY0-1296
PWY-7663
PANTO-PWY
PWY66-400
PWY-6151
PWY0-1319
PWY-5667
PYRIDNUCSYN-PWY
THRESYN-PWY
RIBOSYN2-PWY
PWY-3001
SER-GLYSYN-PWY
NONOXIPENT-PWY
PWY-6471
PWY-5103
PWY-2941
PWY-7198
PWY0-1261
HSERMETANA-PWY
GLUCONEO-PWY
HISTSYN-PWY
THISYN-PWY
PWY-5188
PWY-6168
DTDPRHAMSYN-PWY
PWY0-1241
PRPP-PWY
P42-PWY
PWY66-409
PWY-6969
TCA
HEXITOLDEGSUPER-PWY
PWY-5100
PWY-5104
PWY-6147
PWY-7211
PWY-5101
PWY-6897
0.0
0.5
1.0
1.5
2.0
Weight (%)

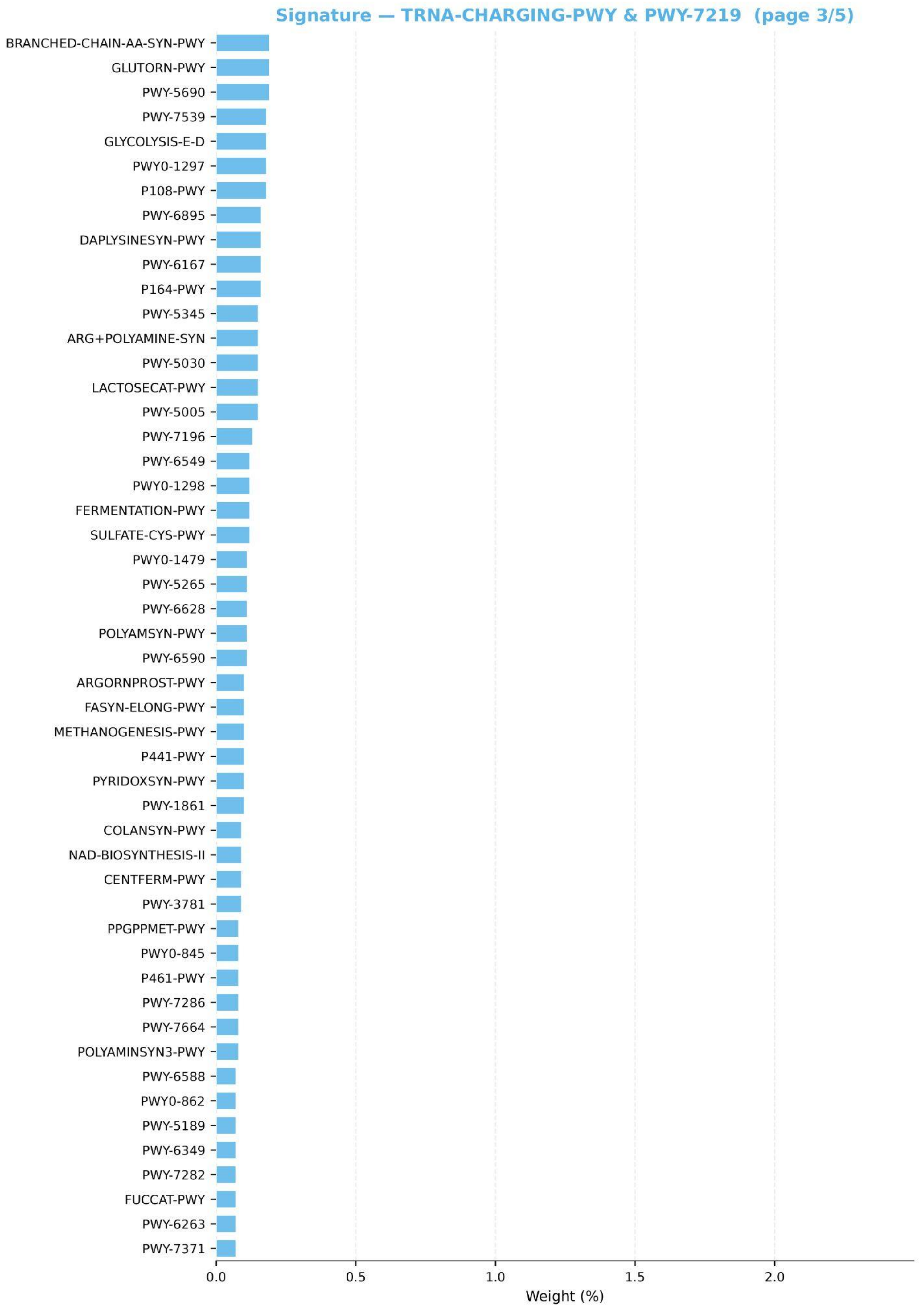

Signature — TRNA-CHARGING-PWY & PWY-7219 (page 3/5)
BRANCHED-CHAIN-AA-SYN-PWY
GLUTORN-PWY
PWY-5690
PWY-7539
GLYCOLYSIS-E-D
PWY0-1297
P108-PWY
PWY-6895
DAPLYSINESYN-PWY
PWY-6167
P164-PWY
PWY-5345
ARG+POLYAMINE-SYN
PWY-5030
LACTOSECAT-PWY
PWY-5005
PWY-7196
PWY-6549
PWY0-1298
FERMENTATION-PWY
SULFATE-CYS-PWY
PWY0-1479
PWY-5265
PWY-6628
POLYAMSYN-PWY
PWY-6590
ARGORNPROST-PWY
FASYN-ELONG-PWY
METHANOGENESIS-PWY
P441-PWY
PYRIDOXSYN-PWY
PWY-1861
COLANSYN-PWY
NAD-BIOSYNTHESIS-II
CENTFERM-PWY
PWY-3781
PPGPPMET-PWY
PWY0-845
P461-PWY
PWY-7286
PWY-7664
POLYAMINSYN3-PWY
PWY-6588
PWY0-862
PWY-5189
PWY-6349
PWY-7282
FUCCAT-PWY
PWY-6263
PWY-7371
0.0
0.5
1.0
1.5
2.0
Weight (%)

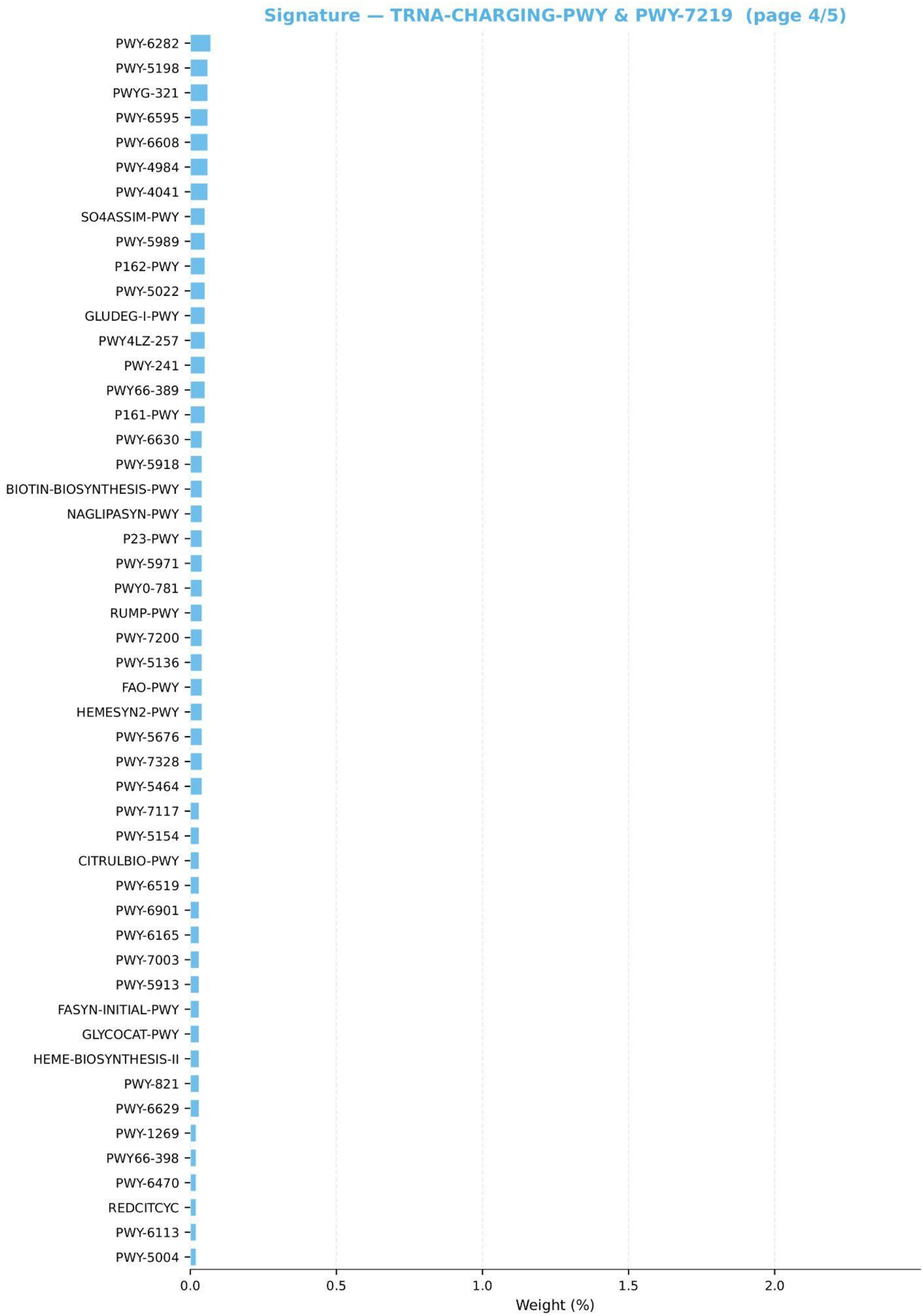

Signature — TRNA-CHARGING-PWY & PWY-7219 (page 4/5)
PWY-6282
PWY-5198
PWYG-321
PWY-6595
PWY-6608
PWY-4984
PWY-4041
SO4ASSIM-PWY
PWY-5989
P162-PWY
PWY-5022
GLUDEG-I-PWY
PWY4LZ-257
PWY-241
PWY66-389
P161-PWY
PWY-6630
PWY-5918
BIOTIN-BIOSYNTHESIS-PWY
NAGLIPASYN-PWY
P23-PWY
PWY-5971
PWY0-781
RUMP-PWY
PWY-7200
PWY-5136
FAO-PWY
HEMESYN2-PWY
PWY-5676
PWY-7328
PWY-5464
PWY-7117
PWY-5154
CITRULBIO-PWY
PWY-6519
PWY-6901
PWY-6165
PWY-7003
PWY-5913
FASYN-INITIAL-PWY
GLYCOCAT-PWY
HEME-BIOSYNTHESIS-II
PWY-821
PWY-6629
PWY-1269
PWY66-398
PWY-6470
REDCITCYC
PWY-6113
PWY-5004
0.0
0.5
1.0
1.5
2.0
Weight (%)

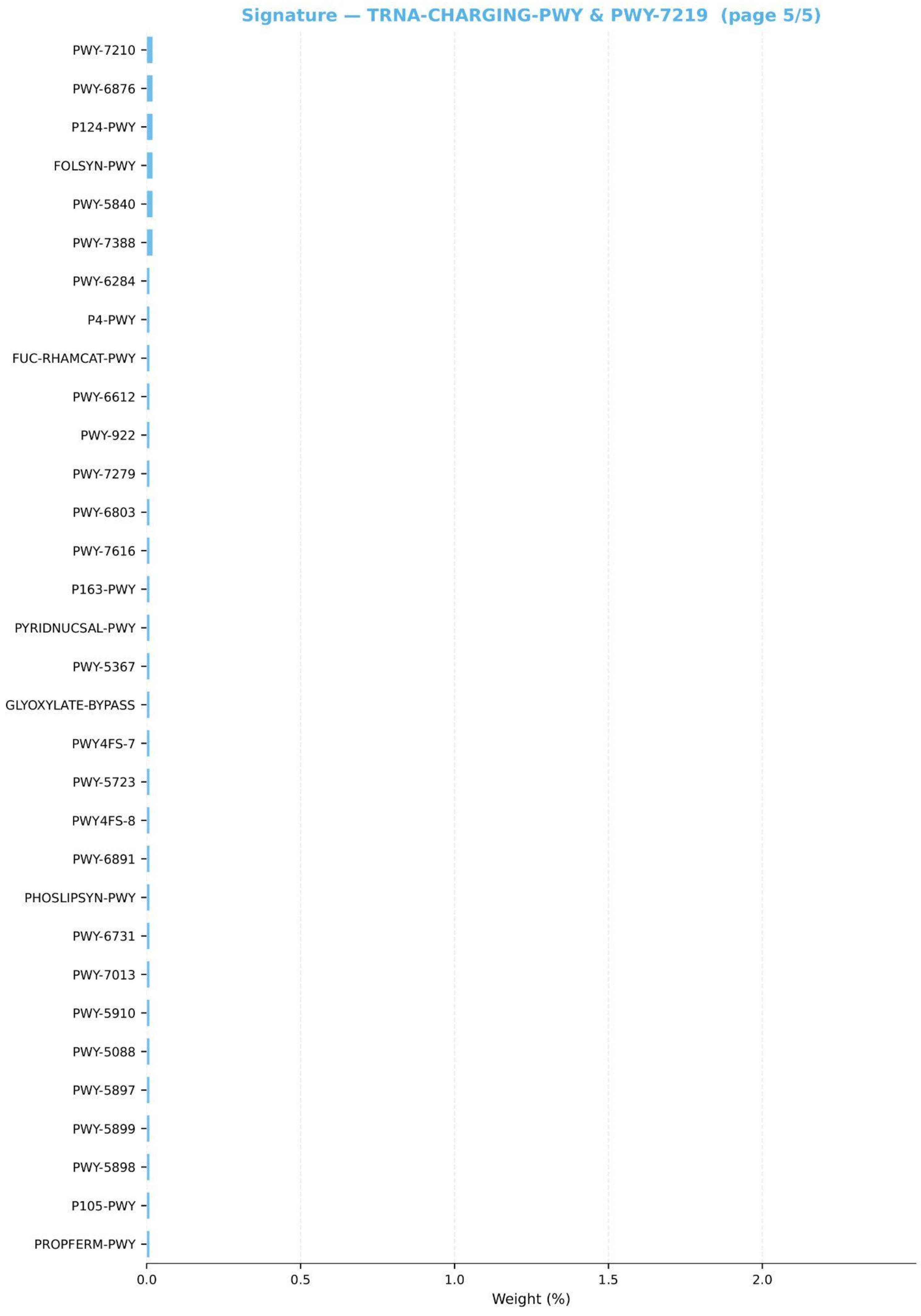

Signature — TRNA-CHARGING-PWY & PWY-7219 (page 5/5)
PWY-7210
PWY-6876
P124-PWY
FOLSYN-PWY
PWY-5840
PWY-7388
PWY-6284
P4-PWY
FUC-RHAMCAT-PWY
PWY-6612
PWY-922
PWY-7279
PWY-6803
PWY-7616
P163-PWY
PYRIDNUCSAL-PWY
PWY-5367
GLYOXYLATE-BYPASS
PWY4FS-7
PWY-5723
PWY4FS-8
PWY-6891
PHOSLIPSYN-PWY
PWY-6731
PWY-7013
PWY-5910
PWY-5088
PWY-5897
PWY-5899
PWY-5898
P105-PWY
PROPFERM-PWY
0.0
0.5
1.0
1.5
2.0
Weight (%)

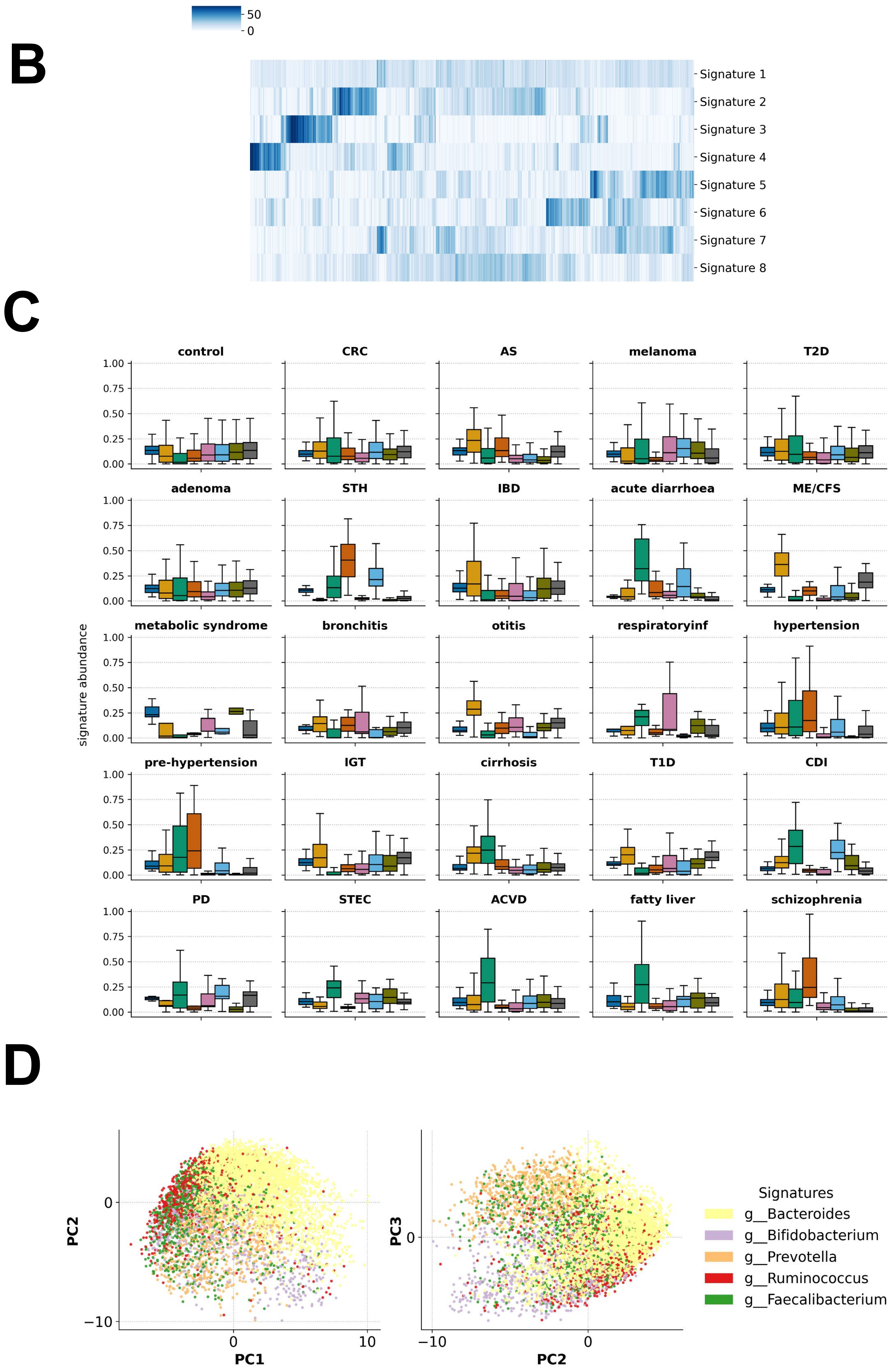


**Supplementary Figure 2. Functional signatures overview.** B. Clustermap showing functional signatures abundance across samples in the training set; C. Functional signatures abundance in different diseases; D. PCoA showing taxonomic NMF-based signatures across samples in the training dataset.

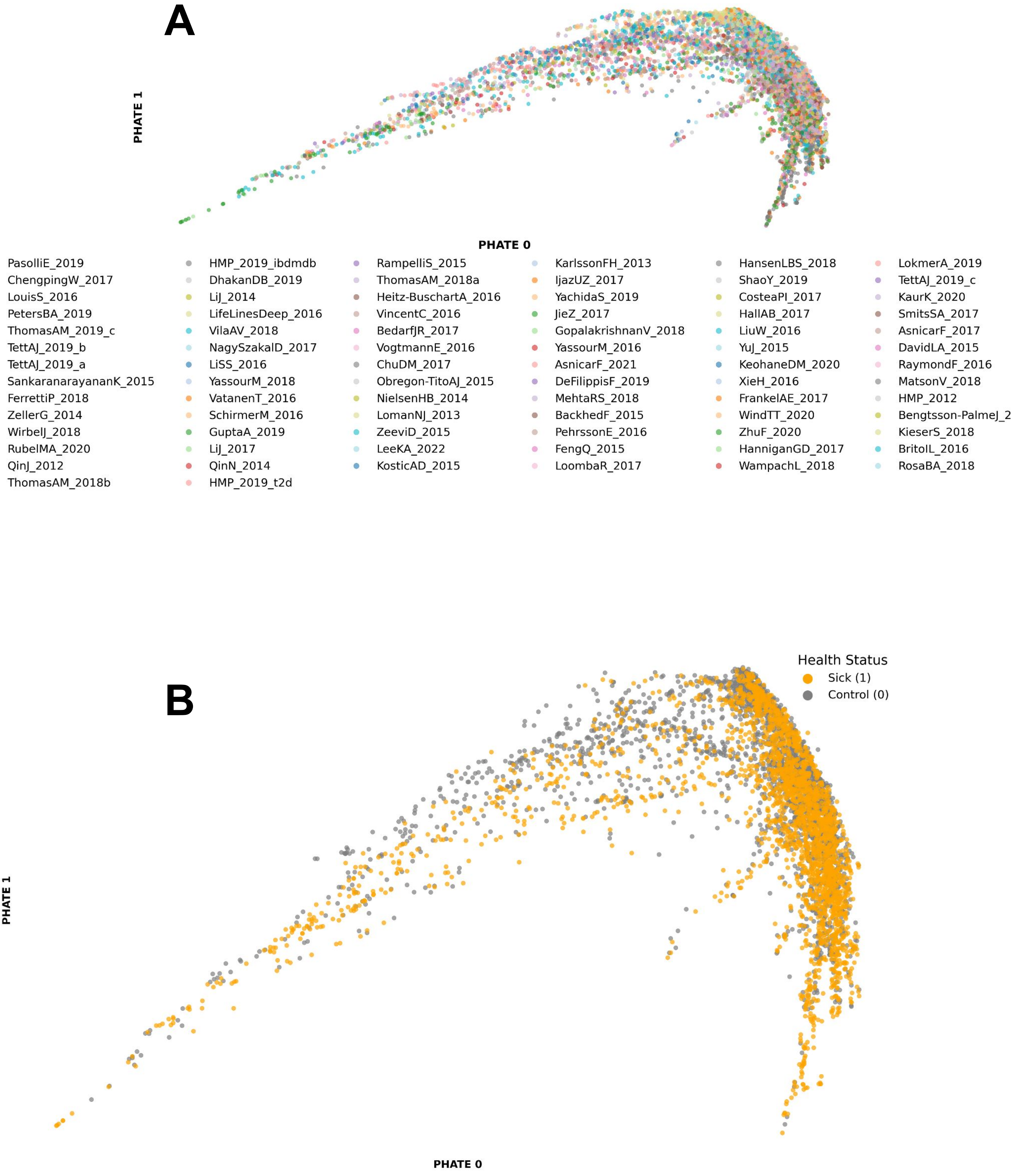


**Supplementary Fig. 3. PHATE visualization of GUT-FORMer learned representations of human gut microbiome samples in the training set.** A. Each point represents a subject; colors indicate dataset origin. B. Each point represents a subject; colors indicate health status, where 1 denotes diseased individuals (any disease) and 0 denotes healthy controls.

**C**

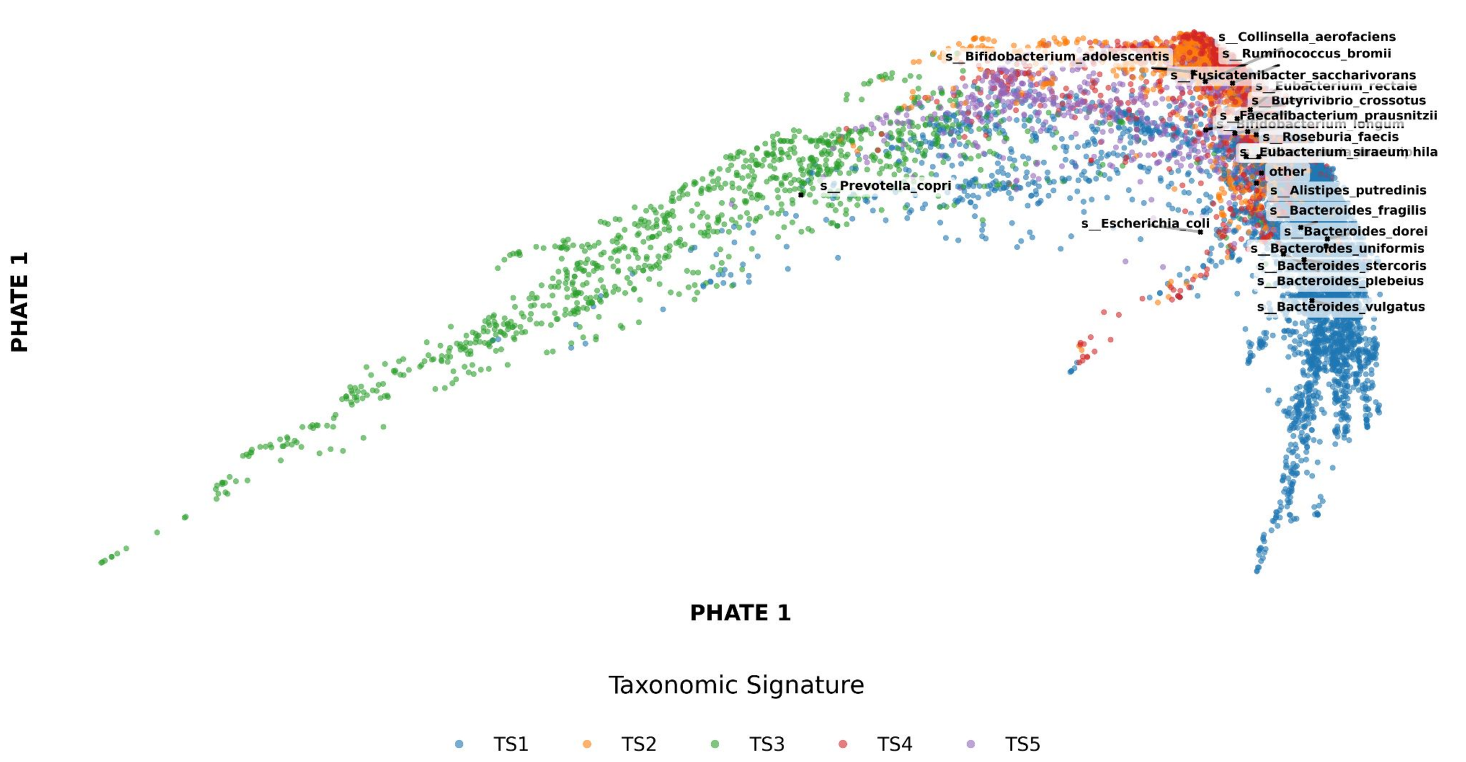


**D**

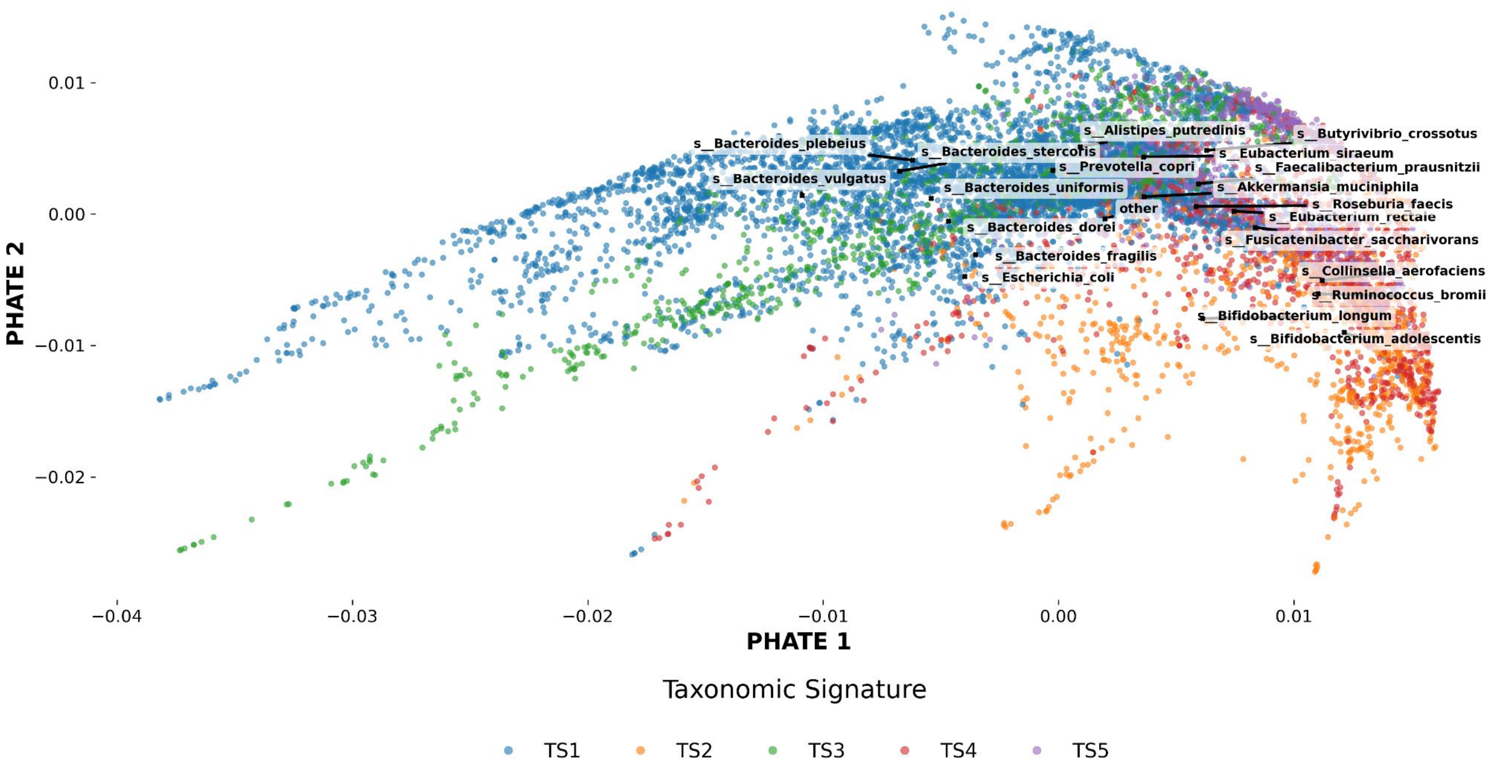


**Supplementary Fig. 3. PHATE visualization of GUT-FORMer learned representations of human gut microbiome samples in the training set.** E. PHATE representation colored by the dominant specie.

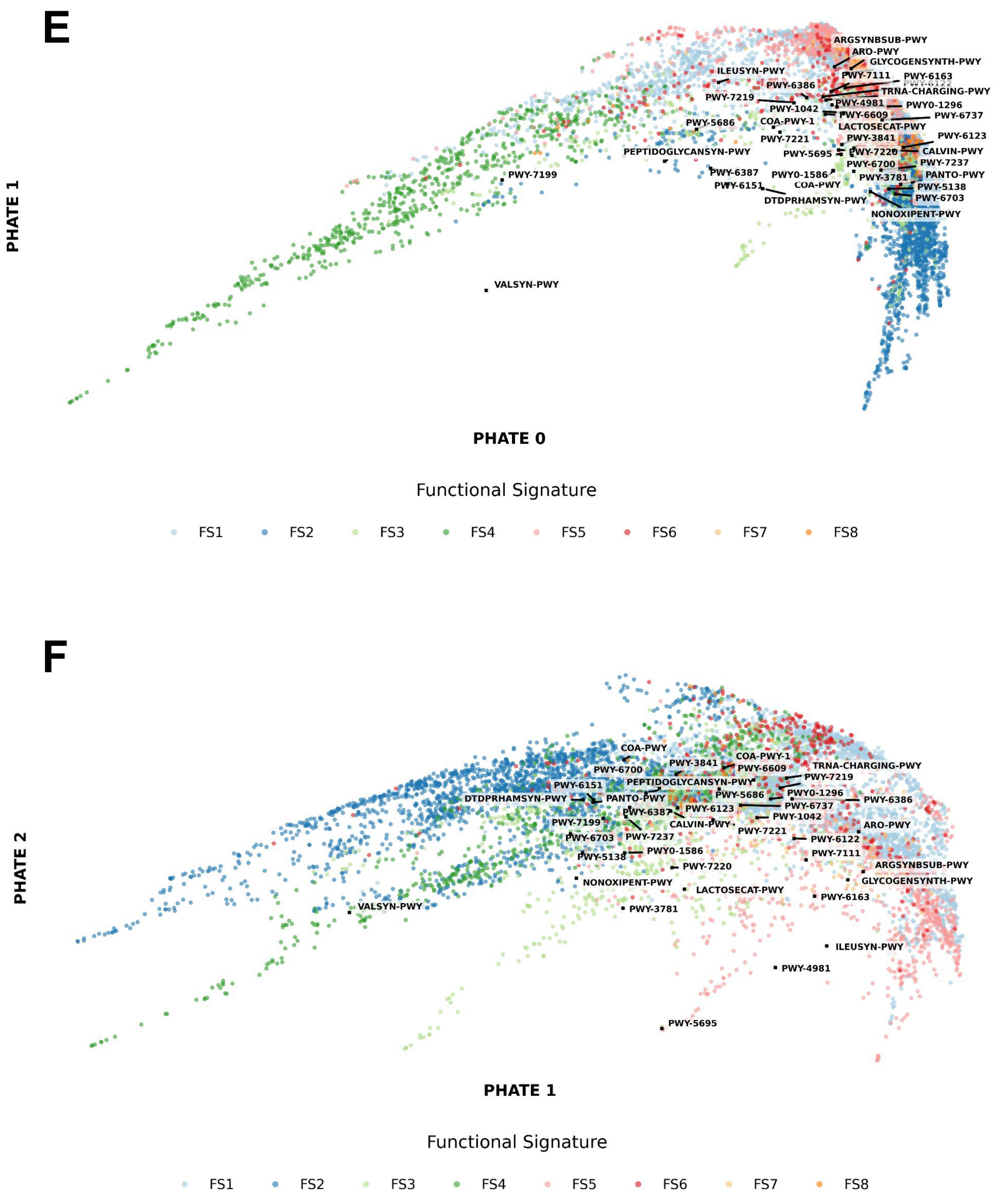

Functional Signature
FS1
FS2
FS3
FS4
FS5
FS6
FS7
FS8

**Supplementary Fig. 3. PHATE visualization of GUT-FORMer learned representations of human gut microbiome samples in the training set.** E. PHATE representation colored by the dominant function.

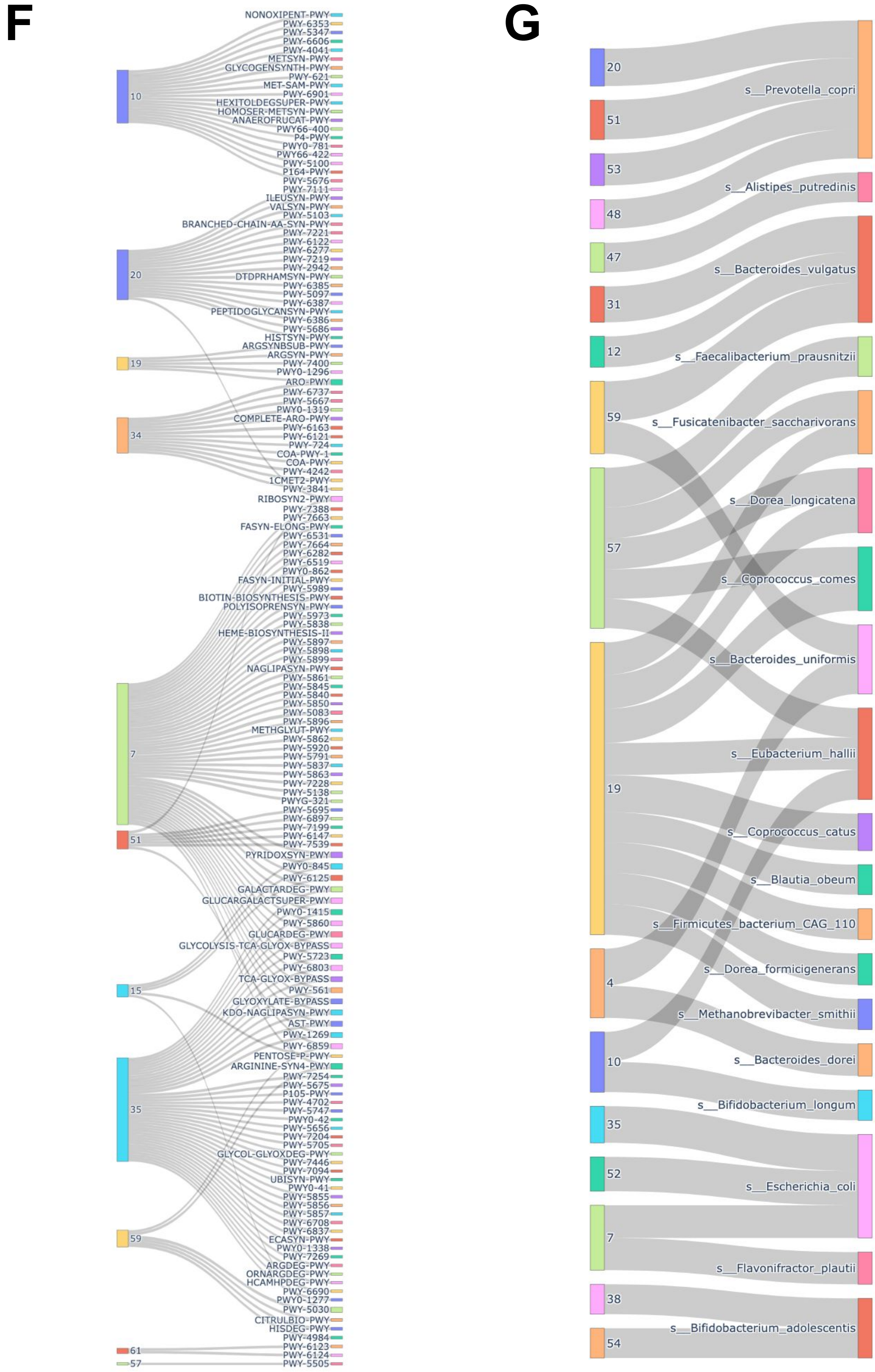


**Supplementary Fig. 3. PHATE visualization of GUT-FORMer learned representations of human gut microbiome samples in the training set.** F. Sankey plot illustrating the relationships between latent dimensions learned by GUT-FORMer and metabolic pathways. G. Sankey plot illustrating the relationships between latent dimensions learned by GUT-FORMer and microbial species. Connections are based on Pearson correlation coefficients, with only correlations > 0.3 retained.

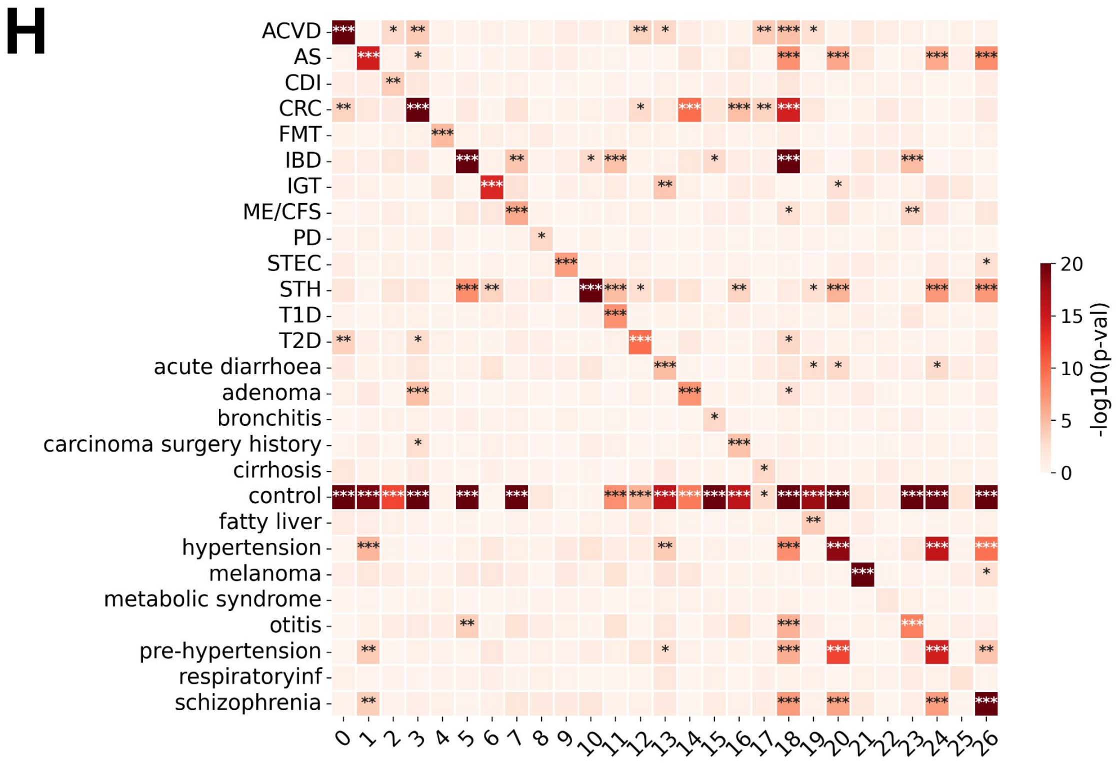


**Supplementary Fig. 3. PHATE visualization of GUT-FORMer learned representations of human gut microbiome samples in the training set.** H. Associations between GUT-FORMer latent dimensions and disease labels defined by FACTM. Asterisks indicate statistical significance (Wilcoxon rank-sum test) for associations between rotated latent dimensions and binary disease status.

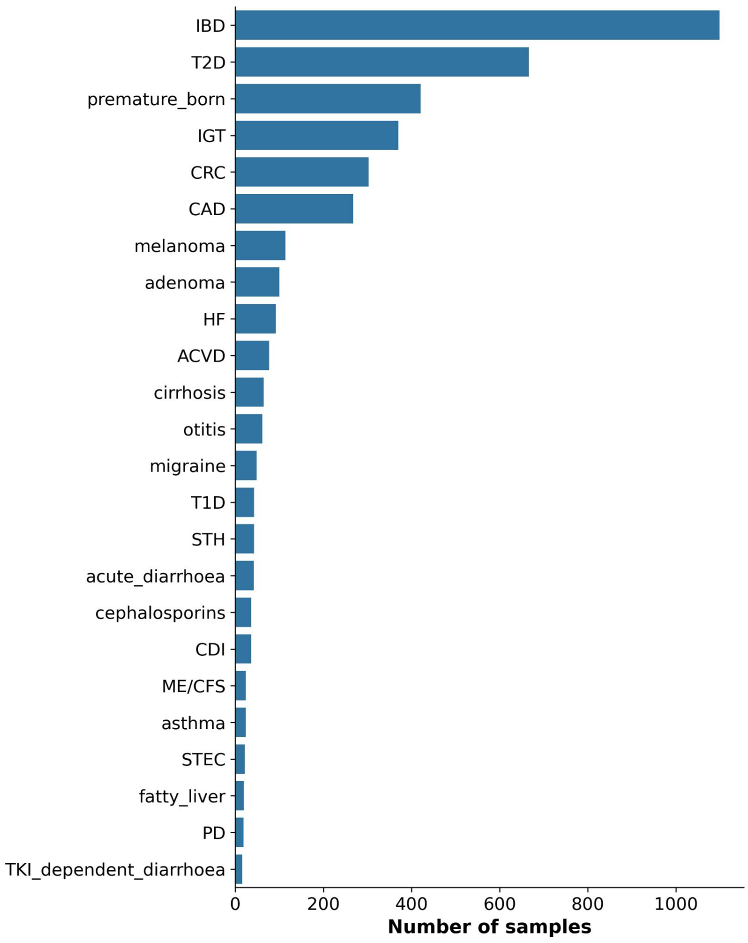


**Supplementary Fig. 4. Distribution of diseases in validation set**

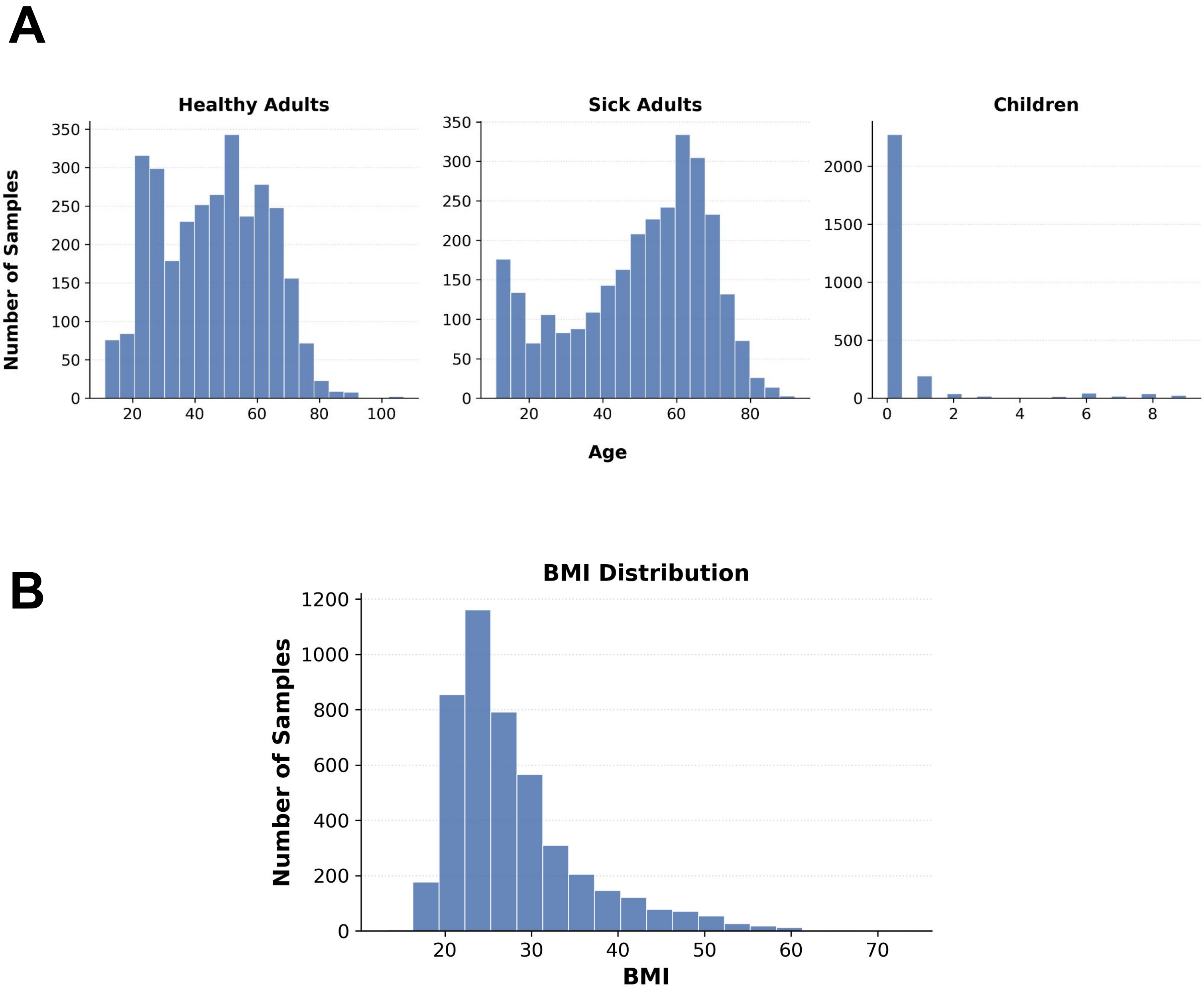


**Supplementary Fig. 5. Regression task data distribution.** A. Age distribution across samples. B. BI distribution across samples (adults only)

A

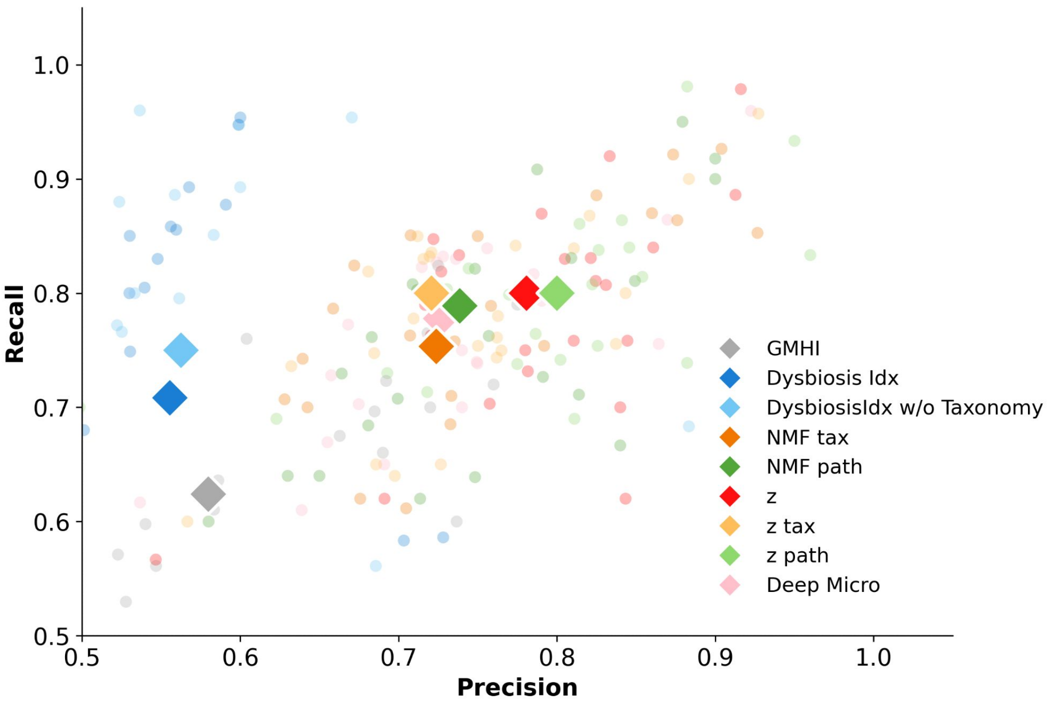


B

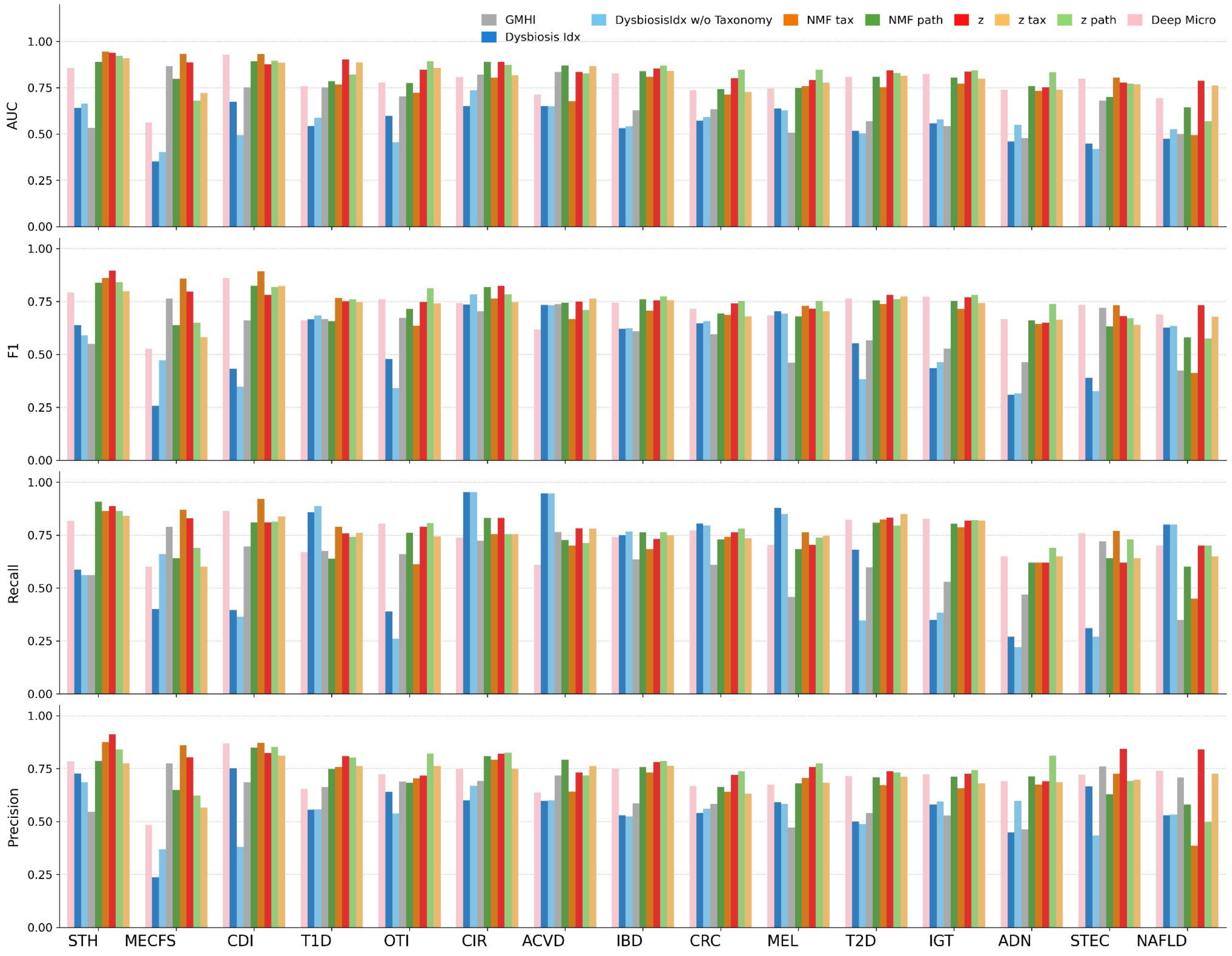


**Supplementary Fig. 5. Benchmarking GUT-FORMer against competing models.** A. Scatterplot showing median Precision and Recall across different models B. Models performance across individual disease classification tasks, where models are trained to distinguish a target disease from healthy..

C

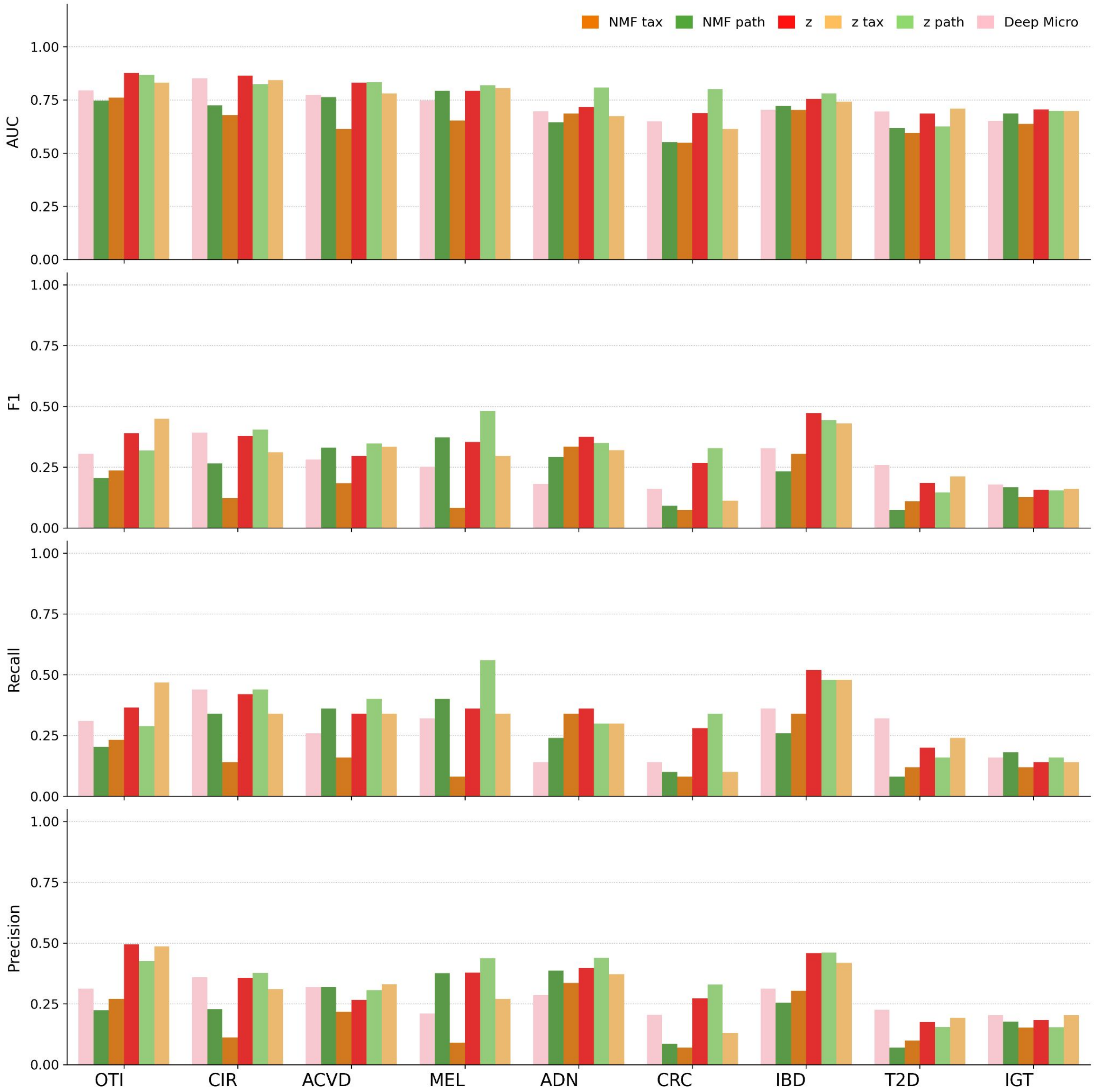


**Supplementary Fig. 6. Benchmarking GUT-FORMer against other models** C. Multiclass classification results for different models across diseases

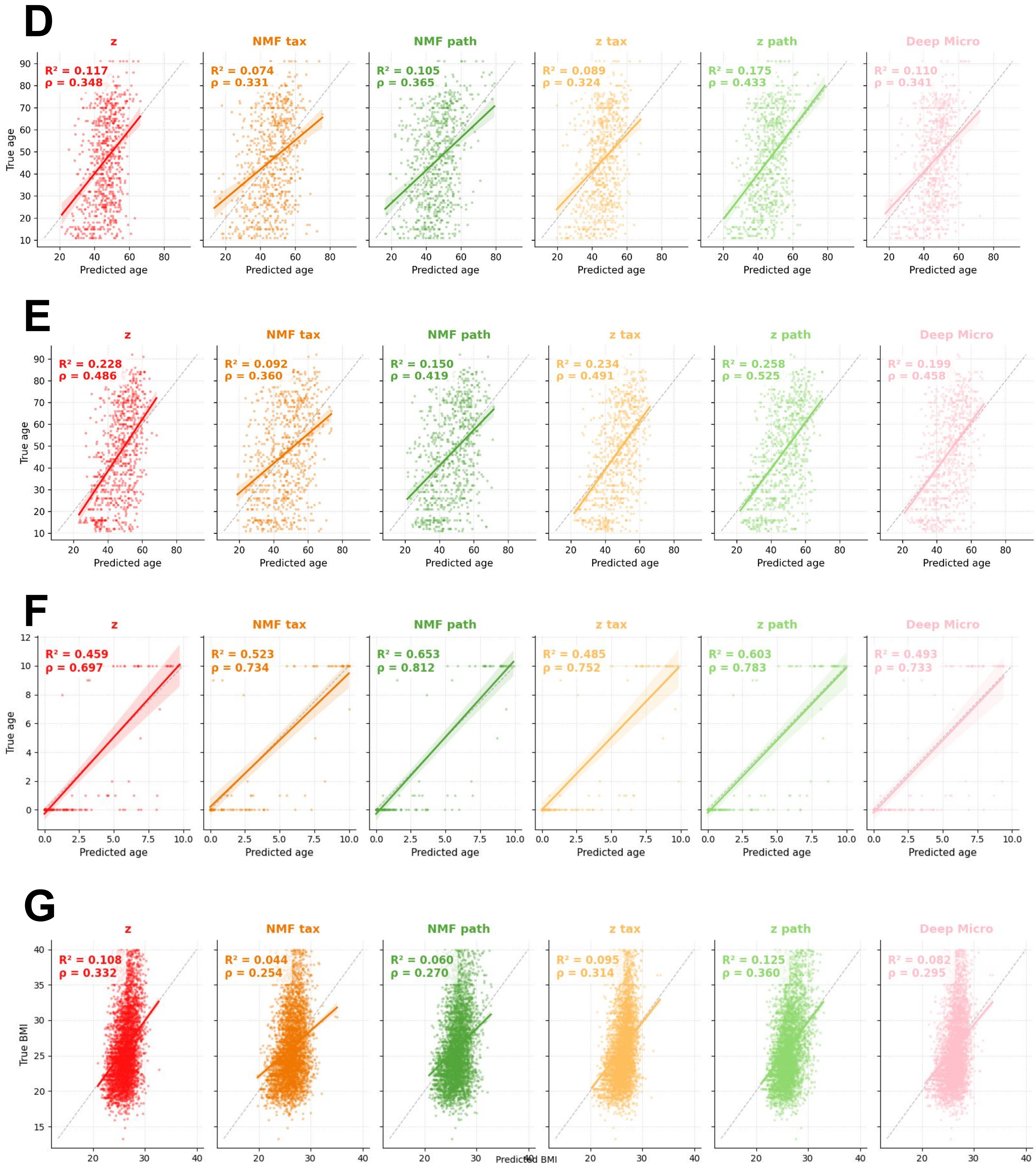


**Supplementary Fig. 6. Benchmarking GUT-FORMer against competing models in regression tasks (age prediction).** D. Age prediction performance across different representation methods in healthy subjects; E. Age prediction performance across different representation methods in diseased subjects; F. Age prediction performance across different representation methods in healthy children. G. BMI prediction performance across different representation methods.

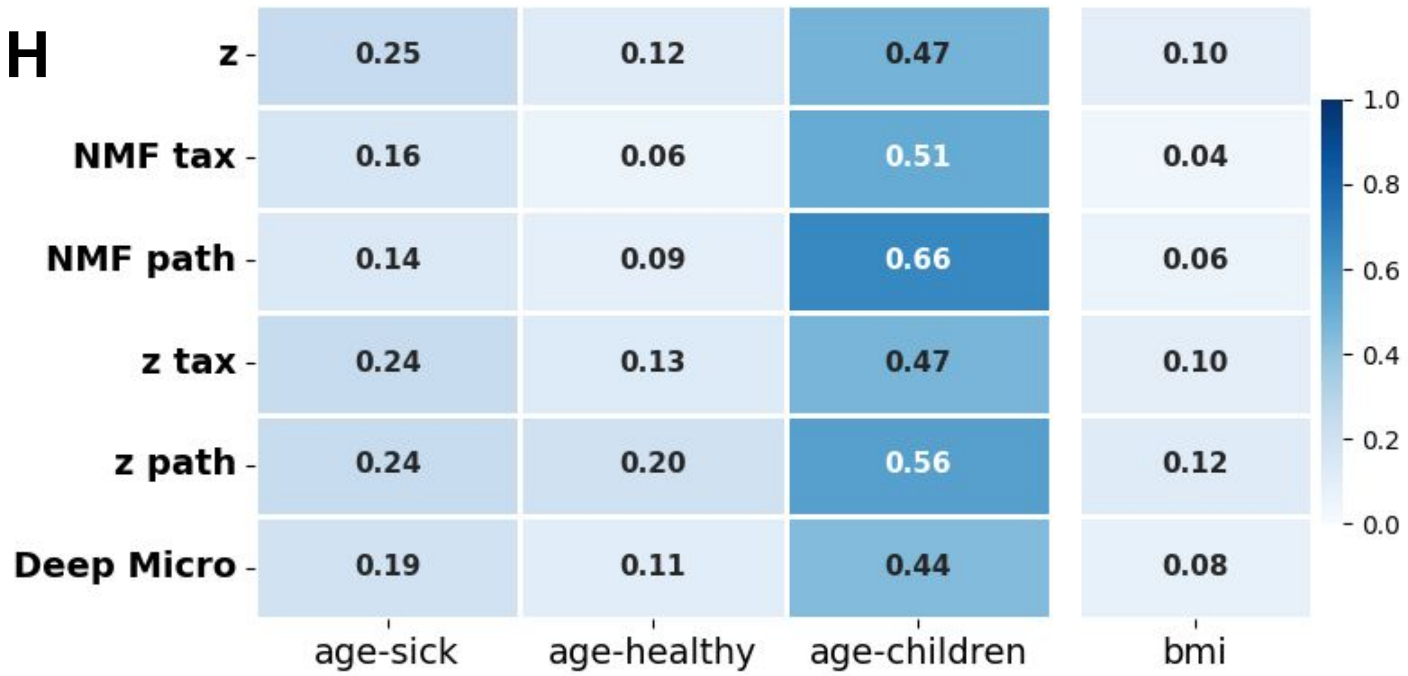


**Supplementary Fig. 6. Benchmarking GUT-FORMer against competing models in regression tasks.** H. R-squared ($R^2$) performance for age and BMI prediction across different models.
